# Inpp5e Is Critical for Photoreceptor Outer Segment Maintenance

**DOI:** 10.1101/2024.08.27.609873

**Authors:** Mohona Gupta, Tylor R. Lewis, Michael W. Stuck, William J. Spencer, Natalia V. Klementieva, Vadim Y. Arshavsky, Gregory J. Pazour

## Abstract

In humans, inositol polyphosphate-5-phosphatase e (INPP5E) mutations cause retinal degeneration as part of Joubert and MORM syndromes and can also cause non-syndromic blindness. In mice, mutations cause a spectrum of brain, kidney, and other anomalies and prevent the formation of photoreceptor outer segments. To further explore the function of Inpp5e in photoreceptors, we generated conditional and inducible knockouts of mouse *Inpp5e* where the gene was deleted either during outer segment formation or after outer segments were fully formed. In both cases, the loss of Inpp5e led to severe defects in photoreceptor outer segment morphology and ultimately photoreceptor cell loss. The primary morphological defect consisted of outer segment shortening and reduction in the number of newly forming discs at the outer segment base. This was accompanied by structural abnormalities of the Golgi apparatus, mislocalized rhodopsin, and an accumulation of extracellular vesicles. In addition, knockout cells showed a reduction in the size and prevalence of the actin network at the site of new disc morphogenesis and the occasional formation of membrane whorls instead of discs in a subset of cells. Together, these data demonstrate that Inpp5e plays a critical role in maintaining the outer segment and the normal process of outer segment renewal depends on the activity of this enzyme.

## Introduction

Primary cilia are evolutionarily conserved microtubule-based organelles that regulate cellular signaling pathways and maintain cellular homeostasis. Defects in cilia cause a large group of diseases known as ciliopathies. The ciliopathies include blindness as the light-sensitive outer segment of rod and cone photoreceptors develops from a primary cilium. To form the outer segment, proteins and lipids synthesized in the inner segment of the cell body are transported into the cilium and the ciliary membrane is remodeled into a stack of flattened discs enriched in visual pigments and signaling molecules. In mice, photoreceptor cells become post mitotic at about post-natal day (P) 3 and the process of forming an outer segment from the primary cilium begins about P9/P10 with outer segments reaching full length at about P25 (LaVail, 1973). Disc formation is driven by the actin cytoskeleton where F-actin polymerization pushes the ciliary membrane out to form a flattened ciliary membrane evagination. As the discs form, the actin filaments depolymerize and in rods, the flattened membrane protrusions are further remodeled to form enclosed discs that are no longer continuous with the ciliary membrane (Spencer et al., 2020). The process of disc formation persists throughout an individual’s life as approximately 10% of the discs are shed from the distal tip daily and are replenished by new discs forming at the base (LaVail, 1973; Young, 1967). Mutations affecting the development or maintenance of the outer segment lead to congenital or degenerative diseases of the retina (Gupta and Pazour, 2023).

Pathogenic variants in *INPP5E* lead to retinal degeneration as part of MORM (Jacoby et al., 2009) and Joubert syndromes (Bielas et al., 2009), and as non-syndromic isolated retinal degeneration (Sangermano et al., 2021). The variants causing MORM syndrome typically affect the CAAX box at the C-terminal end of the protein. This motif is prenylated causing the protein to be peripherally associated with membrane. MORM syndrome impairs INPP5E ciliary localization and is characterized by impaired intellectual development, truncal obesity, micropenis, and retinal dystrophy (Hampshire et al., 2006). Joubert syndrome variants typically affect the enzymatic activity of the protein and yield hypoplasia of the cerebellar vermis with variable presentations of retinal dystrophy and renal anomalies (Bielas et al., 2009; Travaglini et al., 2013; Van De Weghe et al., 2022). The variants associated with isolated retinal degeneration are found outside of the catalytic domain or are subtle substitutions such as Asp to Glu or Arg to His (Sangermano et al., 2021).

Inpp5e was oringinally identified as a Golgi-enriched phosphoinositide phosphatase and proposed to regulate Golgi transport (Kong et al., 2000). It was later found to regulate autophagosome fusion with lysosomes (Hasegawa et al., 2016). Inpp5e activity converts the phosphoinositides PI(3,5)P_2_, PI(4,5)P_2_, and PI(3,4,5)P_3_ into PI(3)P, PI(4)P, and PI(3,4)P_2_, respectively (Kisseleva et al., 2000; Kong et al., 2000). Recently Inpp5e has emerged as an important regulator of ciliary phosphoinositide compostion where it functions to convert PI(4,5)P_2_ to PI(4)P in the ciliary shaft and PI(3,4,5)P_3_ to PI(3,4)P_2_ in the transition zone (Dyson et al., 2017). Under normal conditions, the plasma membrane and ciliary membrane near the ciliary base are enriched in PI(4,5)P_2_ whereas the more distal ciliary membrane has low PI(4,5)P_2_ but elevated PI(4)P due to the action of Inpp5e. This is critically important for the ability of the cilium to regulate Hedgehog signaling (Chavez et al., 2015; Constable et al., 2020; Garcia-Gonzalo et al., 2015) and also regulates removal of cilia through an actin-mediated decapitation of the ciliary tip (Phua et al., 2017). At the transition zone, the lack of Inpp5e activity disrupts the localization of numerous transition zone proteins suggesting that assembly of this structure is dependent on the ratio of PI(3,4,5)P_3_ to PI(3,4)P_2_ (Dyson et al., 2017). The cellular localization of Inpp5e may be regulated, as the loss of the small GTPase Arl16 redistributes Inpp5e from the cilium to the Golgi complex (Dewees et al., 2022).

Germline deletion of *Inpp5e* in mouse causes brain malformations (anencephaly and exencephaly), polydactyly and other skeletal anomalies, cystic kidneys, and bilateral anophthalmia (Jacoby et al., 2009). Deletion of *Inpp5e* in the retina with *Six3-Cre* blocked the formation of discs from the ciliary membrane and caused rapid retinal degeneration (Sharif et al., 2021). This Cre is expressed embryonically in the retina prior to cells becoming post-mitotic and forming primary cilia (Furuta et al., 2000) indicating that Inpp5e is critical to early stages of photoreceptor outer segment development. To overcome challenges posed by embryonic lethality and to uncover Inpp5e function in later stages of outer segment formation and in photoreceptor maintenance, we utilized two distinct conditional mutant mouse models. In the first model, an *Inpp5e* floxed allele was deleted by rod-specific *iCre75* during photoreceptor outer segment maturation. In the second model, the *Inpp5e* floxed allele was deleted by the tamoxifen inducible rod-specific *Pde6g^CreERT2^*, which allowed us to analyze Inpp5e function in mature photoreceptors after outer segment morphogenesis was complete. In both cases, the loss of Inpp5e led to rapid photoreceptor degeneration. The degeneration was accompanied by shortening of the outer segments, malformations of the Golgi, accumulation of rhodopsin in the cell body, abnormal actin cytoskeleton, and the presence of abundant extracellular vesicles around photoreceptor somas. These findings suggest Inpp5e plays a critical role in outer segment maintenance.

## Results

### Rod-specific Inpp5e knockout leads to rapid vision loss

To elucidate the role of Inpp5e in photoreceptors, we deleted a floxed allele of *Inpp5e* (Dyson et al., 2017) using two Cre recombinase lines. The first, *iCre75*, utilizes a rod opsin promoter to selectively express Cre recombinase in rod photoreceptor cells. In this line, Cre expression begins at about postnatal day 7 (P7) (Li et al., 2005). Outer segment elongation starts at about P9/P10 (LaVail, 1973; Obata and Usukura, 1992; Pearring et al., 2013), thus it is expected that pathology would start during the elongation phase due to the time required for the gene to be deleted and the gene products decay. The second, *Pde6g^CreERT2^* expresses a tamoxifen-inducible Cre recombinase under the control of the endogenous rod-specific *Pde6g* promoter allowing for *Inpp5e* ablation in mature rods. The *Inpp5e* floxed allele contains loxP sites flanking the catalytic domain and is expected to produce a null allele after deletion (Dyson et al., 2017).

Electroretinography (ERG) was used to gauge photoreceptor function. In the *iCre75* line, experimental (*Inpp5e^flox/flox^/iCre75*) animals showed a clear reduction in retinal function at P21 (Fig 1A). The retinal dysfunction was evident in both the initial rod photoreceptor response to light (scotopic a wave) and the downstream responses (scotopic b wave) when compared to control (*Inpp5e^flox/+^/iCre75*) animals. Scotopic a and b waves were nearly absent at P30, although a residual signal was observed that could be driven by cells where the Cre was not effective or in cells where the outer segments have not yet fully degenerated. Cone-driven responses (photopic b wave) were not significantly affected at these points. This is expected as we deleted the *Inpp5e* gene in rod photoreceptors and the time points examined are too early for secondary cone loss.

**Figure 1.**
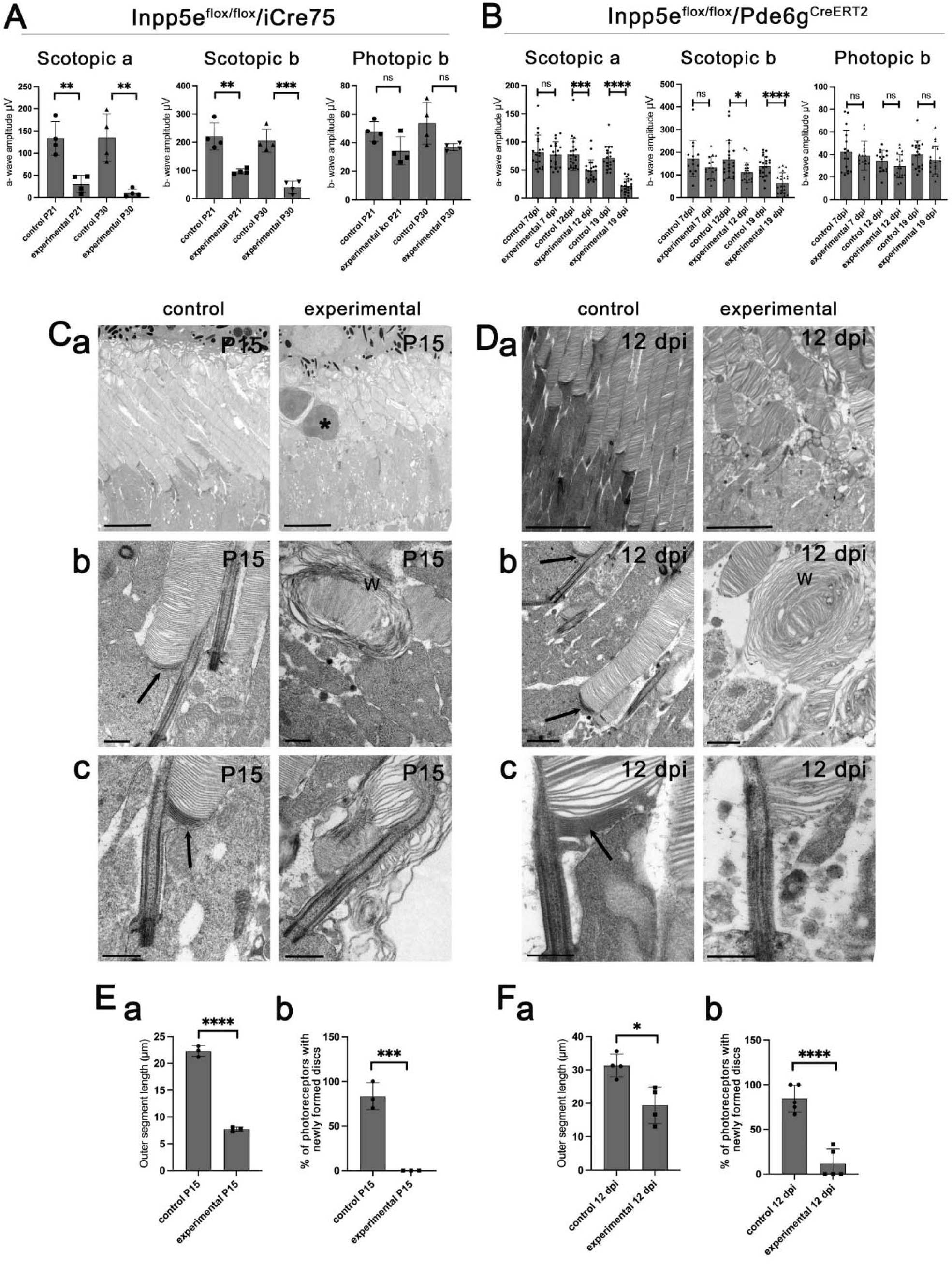
Loss of Inpp5e leads to rapid vision loss. (A) Electroretinograms of *Inpp5e^flox/flox^* (control) and *Inpp5e^flox/flox^/iCre75* (experimental) littermates examined at P21 and P30. Each point represents the average response from one animal. **p<0.01, ***p<0.001, ns not significant by unpaired t-test. (B) Electroretinograms of *Inpp5e^flox/flox^/Pde6g^CreERT2^* littermates treated with vehicle (control) or tamoxifen (experimental) and examined 7, 12, and 19 days post last injection (dpi). Each point represents the average response from one animal. *p<0.05, ***p<0.001, ****p<0.0001, ns not significant by unpaired t-test. (C) TEM images of *Inpp5e^flox/flox^* (control) and *Inpp5e^flox/flox^/iCre75* (experimental) littermates examined at P15. Scale bar is 5 µm in Ca, 0.5 µm in Cb and Cc. The asterisk marks the nuclei of a dead photoreceptor, w marks whorls of membrane, arrows point to darkened membrane at the base of outer segments. (D) TEM of *Inpp5e^flox/flox^/Pde6g^CreERT2^* littermates treated with vehicle (control) or tamoxifen (experimental) examined 12 days post last injection (dpi). Scale bar is 5 µm in Da, 0.5 µm in Db, and Dc. w marks membrane whorls, arrows point to darkened membrane at the base of outer segments. (Ea and Fa) Average outer segment length as measured in TEM of *iCre75* and *Pde6g^CreERT2^* animals as P15 and 12 dpi respectively. Each point is the average from one animal. *p<0.05, ****p<0.0001 by unpaired t test. (Eb and Fb) Percent of photoreceptors with newly formed discs in TEMs of *iCre75* and *Pde6g^CreERT2^* animals as P15 and 12 dpi respectively. Each point is the average from one animal. In all panels, error bars represent standard deviations. Significance ***p<0.001, ****p<0.0001 by unpaired t-test.

For analysis of Inpp5e in mature photoreceptors, *Inpp5e^flox/flox^/Pde6g^CreERT2^*mice were treated with tamoxifen (experimental) or vehicle (control). Initially, we examined retinas at 30 days post last injection (dpi), but at this point the retinas were highly degenerated as observed by a severely thinned outer nuclear layer and little to no detectable outer segment layer (Fig S1). To identify points early in degeneration where the effects on the retina are more likely directly caused by the loss of Inpp5e rather than the indirect effects of cell death, we carried out ERGs to monitor visual function at 7, 12 and 19 dpi (Fig 1B). At 7 dpi, ERG responses of experimental mice were not different from controls, but by 12 dpi, scotopic a- and b-waves were reduced, and decreased further at 19 dpi. No significant difference in cone-driven responses (photopic b-waves) were observed at any time point, consistent with the *Inpp5e* knockout being rod specific.

### Loss of Inpp5e blocks new disc formation

Transmission electron microscopy (TEM) was used to assess photoreceptor structure. For *iCre75* deletion, dysfunction as measured by ERG was already evident at P21 (Fig 1A), prompting tissue collection at P15 in order to survey the pathology earlier in disease progression. For *Pde6g^CreERT2^*animals, we collected tissue at 12 dpi, as this was the point when dysfunction was first detected by ERG. The pathology detected by TEM was similar in retinas collected from both lines (Fig 1C, 1D). Both models, most experimental outer segments were irregularly shaped and shortened to about ½ as long as the control outer segments (Fig 1Ca, 1Da, 1Ea, 1Fa). Structural abnormalities included malformed discs and occasional membrane whorls (Fig 1Cb, 1Db). In the *iCre75* model, we observed occasional nuclei in the outer segment layer suggesting photoreceptor cell death was occurring (Fig 1Ca). To examine the formation of new discs, we contrasted tissue with tannic acid/uranyl acetate. Due to its low membrane permeability, tannic acid labels cell surface-exposed membranes more strongly than internal membranes. Hence, this technique labels the membranes of newly forming discs exposed to the extracellular environment more darkly than membranes of fully formed discs enclosed inside the outer segment (Ding et al., 2015) (Fig 1Cc, 1Dc). In both lines, darkly stained discs at the base of the outer segment were observed in ∼75% of cells in control animals but they were almost never seen in experimental animals (Fig 1Eb, 1Fb). This observation suggests that new disc formation does not occur when Inpp5e is missing.

### Inpp5e knockouts mislocalize rhodopsin

In addition to the defects observed in the outer segments, the inner segment region of experimental animals from both lines contained large amounts of extracellular vesicles (Fig 1Ca, 1Da, 2A) similar to those in models with rhodopsin mislocalization [reviewed in (Spencer, 2023)]. To investigate whether rhodopsin mistrafficking contributes to the formation of extracellular vesicles when Inpp5e is missing, we conducted rhodopsin immunogold electron microscopy. The vesicles labeled strongly with anti-rhodopsin antibodies consistent with them containing rhodopsin (Fig 2B).

**Figure 2.**
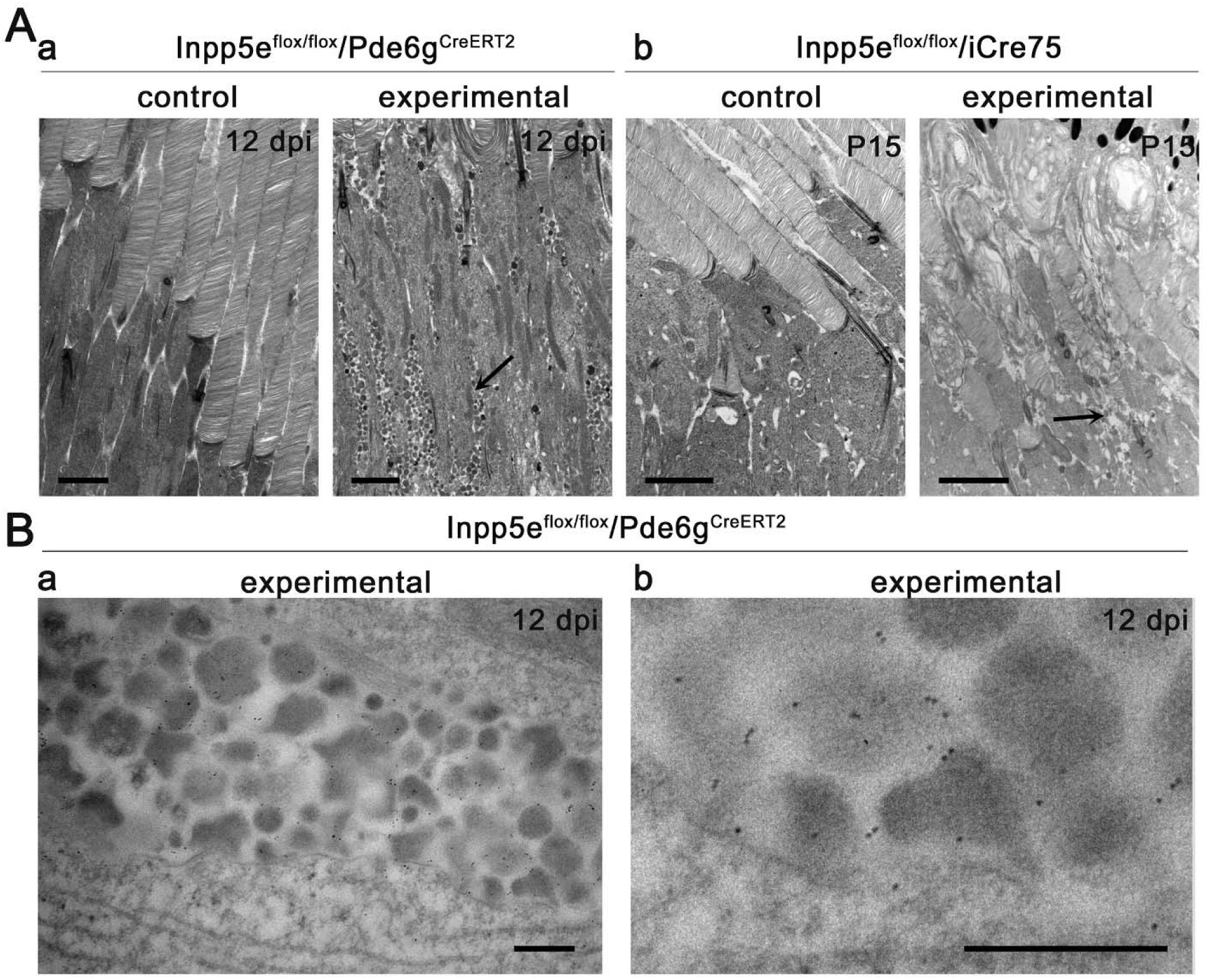
Loss of Inpp5e causes loss of outer segment integrity and extracellular vesicle accumulation. (Aa) TEM of *Inpp5e^flox/flox^/Pde6g^CreERT2^* littermates treated with vehicle (control) or tamoxifen (experimental) examined 12 days post last injection (dpi). (Ab) TEM of *Inpp5e^flox/flox^/iCre75* littermates examined at postnatal day 15 (P15). Images of inner segments reveal accumulation of vesicles (arrow) within the inner segment region of photoreceptors in the experimental group, which is absent in the control group. Scale bar: 2 µm. (B) Immunogold EM shows rhodopsin localization in the extracellular vesicles in the experimental group. The gold particles have been labeled in fuchsia in top row (Ba, Bb). Scale bar: 300 nm.

Rhodopsin, the visual pigment in rod photoreceptors, is synthesized in the inner segment and transported to the outer segment where it is highly concentrated. Accordingly, wild type mice typically display weak rhodopsin signal in the inner segments and strong signal in the outer segments. Retinal degeneration-causing mutations often misaccumulate rhodopsin in the inner segment, nuclear layer and synaptic regions. In the *Pde6g^CreERT2^* model, rhodopsin mislocalization was negligible at 7 dpi but became prominent by 12 dpi (Fig 3). *Inpp5e/iCre75* experimental animals showed rhodopsin accumulation throughout the entire photoreceptor cell body as early as P15, which was further increased at P21 (Fig S2).

**Figure 3.**
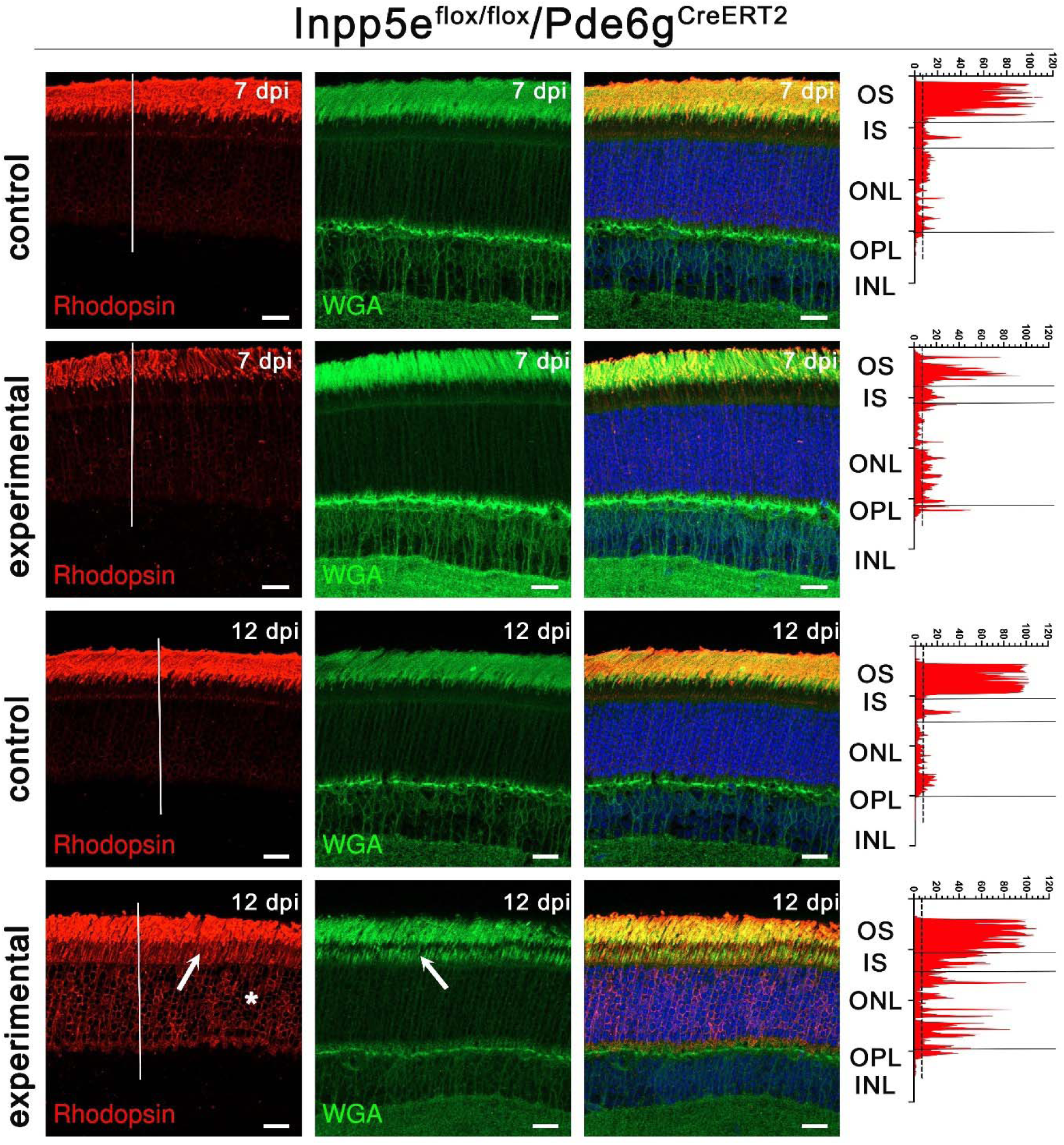
Loss of Inpp5e causes rhodopsin mislocalization. Confocal images of retinal sections (agarose embedded) of *Inpp5e^flox/flox^/Pde6g^CreERT2^* littermates treated with vehicle (control) or tamoxifen (experimental) at 7 and 12 days post last injection (dpi) immunostained with anti-rhodopsin clone 4D2 (red) and wheat germ agglutinin (WGA, green). Scale bars: 20 µm. Each image is a maximum intensity projection of 20 images taken at 0.7-µm intervals. Rhodopsin intensity along the white line is shown on the right side of the images. Arrows point to mislocalized rhodopsin or WGA in the inner segment. * points to rhodopsin in the nuclear layer. OS, outer segment; IS, inner segment; ONL, outer nuclear layer; OPL, outer plexiform layer; INL, inner nuclear layer.

### Golgi structure is disrupted by loss of Inpp5e

Wheat germ agglutinin (WGA) binds to glycosylated proteins, such as rhodopsin, and normally labels outer but not inner segments. In the *Pde6g^CreERT2^* animals, WGA staining was fairly normal at 7 dpi, but was significantly increased in the inner segments and the outer nuclear layer by 12 dpi (Fig 3). In *iCre75* experimental animals, WGA staining was slightly elevated in the inner segment layer by P15 and became prominent by P21 (Fig S2). WGA staining only partially overlapped with rhodopsin staining indicating that it is not solely due to rhodopsin mislocalization. The strongest WGA staining in the inner segment was seen just above the nuclear layer where Golgi complexes are located. To determine if the Golgi structure was affected by the loss of Inpp5e, we stained retinas with WGA and the cis-medial Golgi complex marker giantin (Nozawa et al., 2002). In control animals, giantin prominently labeled elongated structures in the expected Golgi location (Fig 4Aa). In 12 dpi experimental animals, the intensity of giantin label was slightly reduced and the structures observed were more rounded than in control (Fig 4A). TEM of control retinas showed an elongated Golgi structure typical of rod cells whereas the Golgi structures in experimental retinas were harder to identify and the ones observed were typically more swollen (Fig 4B), consistent with light microscopy observations (Fig 4A).

**Figure 4.**
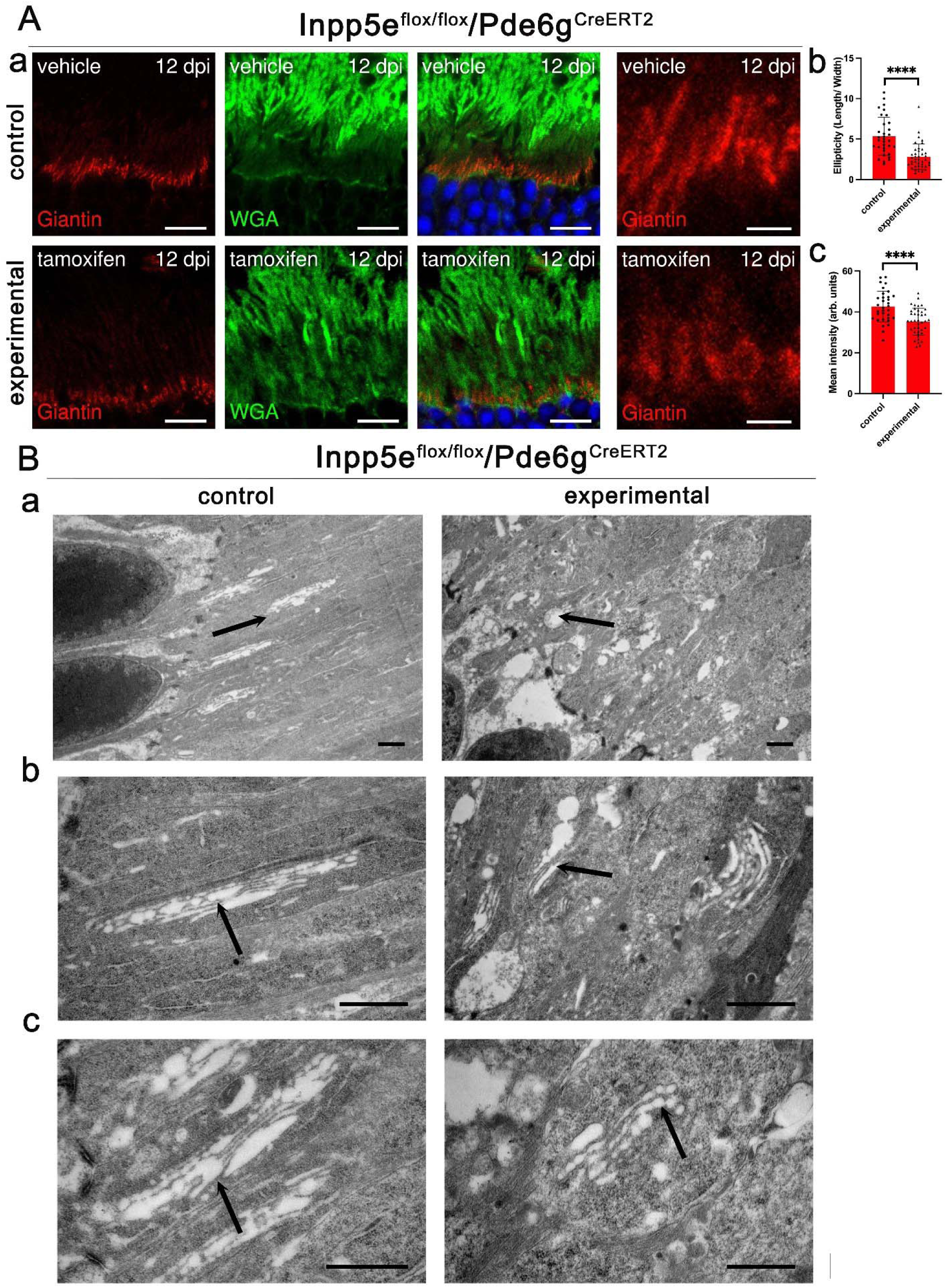
Loss of Inpp5e affects Golgi structure. (Aa) Confocal images of retinal sections (cryosections) of *Inpp5e^flox/flox^/Pde6g^CreERT2^* littermates treated with vehicle (control) or tamoxifen (experimental) at 12 days post last injection (dpi) immunostained with anti-Giantin (red) and wheat germ agglutinin (WGA, green). Scale bar: 10 µm. Each image is a single plane of a z-stack. (Ab) Quantification of ellipticity. Each dot represents the length/width of a Golgi structure stained with giantin. ****p<0.0001 by unpaired t-test. (Ac) Intensity of giantin label per Golgi structure. ****p<0.0001 by unpaired t-test. (B) TEMs showing the inner segment above the limiting membrane where Golgi structures are located. Arrows point to Golgi stacks. Scale bar: 1 µm in all images.

To analyze the extent of protein mislocalization from outer segments in the absence of Inpp5e, we surveyed the subcellular distribution of integral membrane proteins peripherin-2 (Prph2), retinal outer segment membrane protein 1 (Rom1), and prominin-1 (Prom1) (Fig 5, S3). Prph2 and Rom1 support disc rim formation (Stuck et al., 2016) and are normally localized throughout the outer segment. Prom1 is important for disc morphogenesis and is normally found at the base of the outer segment (Yang et al., 2008). We did not observe any notable changes in the distribution of Prph2 or Rom1 in either *Inpp5e* knockout model (Fig 5A, 5B, S3A, S3B). In contrast, there was a minor inner segment accumulation of Prom1 (Fig 5C, S3C). A possible explanation for the different effects of the Inpp5e loss on these proteins’ distribution may lie in their distinct trafficking pathways. Prph2 and Rom1 are delivered to the outer segment directly from the endoplasmic reticulum (Tian et al., 2014), whereas rhodopsin, and potentially Prom1, are processed through the Golgi apparatus before reaching their final destination (Campos et al., 2011; Papermaster et al., 1986; Sung and Tai, 2000). This difference in trafficking routes could account for the accumulation of Prom1 in the inner segment of our *Inpp5e* knockout models. Our findings are similar to those observed in Sharif *et al*. except that the deletion of *Inpp5e* by *Six3- Cre* did not yield a notable increase in inner segment Prom1 (Sharif et al., 2021).

**Figure 5.**
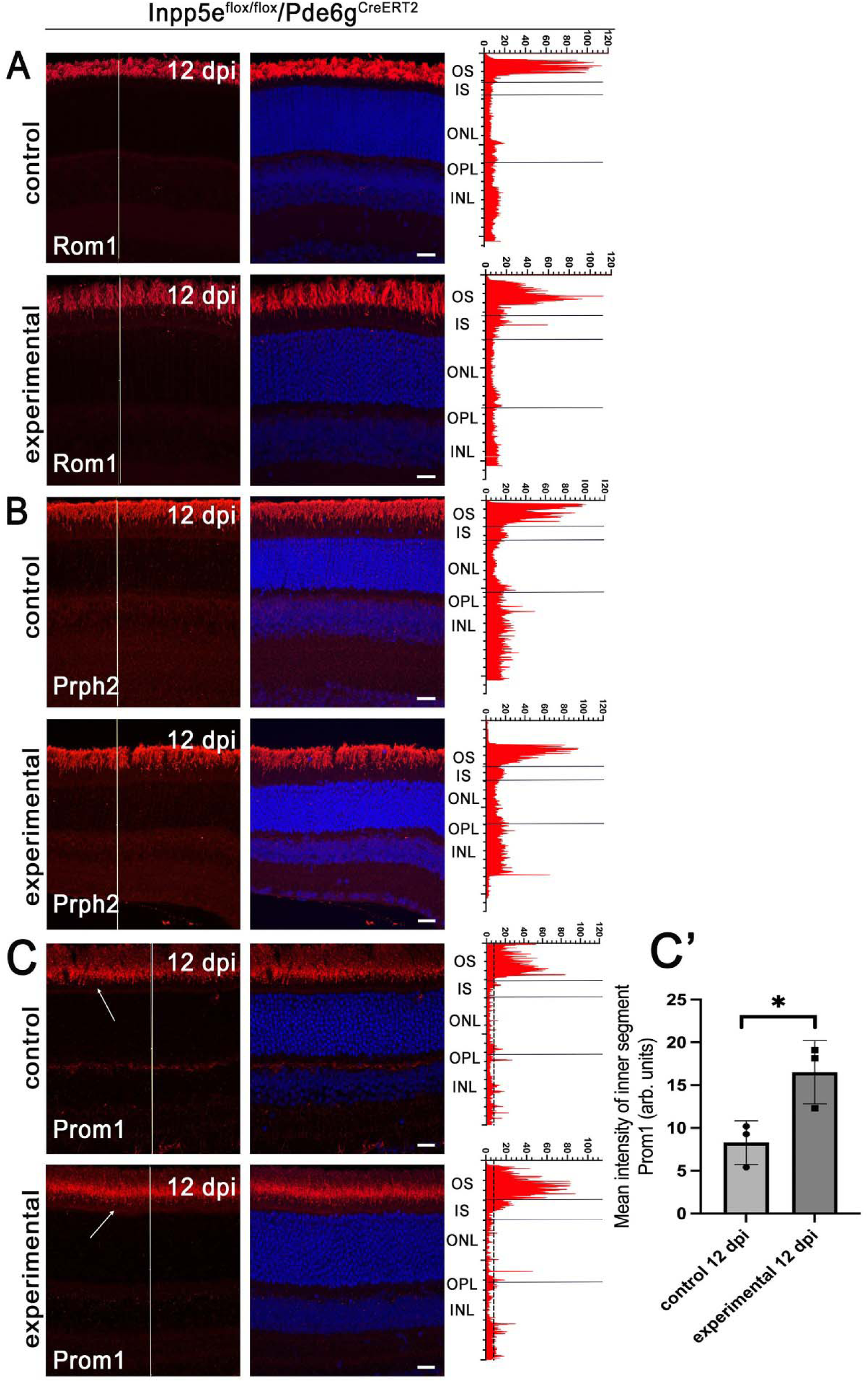
Loss of Inpp5e alters distribution of Prom1. (A-C) Confocal images of retinal sections (agarose embedded) of vehicle (control) and tamoxifen (experimental) treated *Inpp5e^flox/flox^/Pde6g^CreERT2^* littermates at 12 days post last injection (dpi). Sections were stained with DAPI (blue) and for Rom1 (A), Prph2 (B), and Prom1 (C) in red. Note increased Prom1 in the inner segments (arrow) of the experimental animals compared to controls. Scale bar: 20 µm. Each image is a maximum intensity projection of 20 images taken at 0.7-µm intervals. The intensity of the red channel along the white line is shown on the right side of the images. OS, outer segment; IS, inner segment; ONL, outer nuclear layer; OPL, outer plexiform layer; INL, inner nuclear layer. (C’) Quantification of Prom1 fluorescence intensity in the inner segments of control and experimental retinas (n=3, *p<0.05, by unpaired t- test). Each point represents an individual measurement from one animal, with mean ± SD shown.

We also analyzed the distribution of the SNARE protein syntaxin-3 (Stx3), which is normally excluded from the outer segment but present in the inner segment and synaptic terminal (Datta et al., 2015). We found no difference in Stx3 distribution between control and *Inpp5e* knockout rods (Fig S4). Stx3 accumulated in the outer segment in models of retinal degeneration caused by defects in the BBSome (Datta et al., 2015; Dilan et al., 2018; Hsu et al., 2017), but not in *Ift20* and *Ift172* knockouts (Lewis et al., 2024). The lack of Stx3 accumulation in *Inpp5e* knockouts could indicate that *Inpp5e* loss does not affect the trafficking of Stx3. Alternatively, the failure of new disc assembly could close the pathway for Stx3 entry into the cilium, thus indirectly preventing its accumulation.

### Loss of Inpp5e disrupts the actin cytoskeleton

The occasional membrane whorls emanating from the photoreceptor cilium in a subset of *Pde6g^CreERT2^/Inpp5e* mice (Fig 1Cb, 1Db) resembles the overgrown discs observed in mice with disrupted actin network regulators (Spencer et al., 2019; Spencer et al., 2023). To determine if the actin cytoskeleton is disrupted by the loss of Inpp5e, we stained retinas with fluorescent phalloidin, which labels F-actin (Fig6, S5). In control retinas, phalloidin strongly stained the inner segment region with the most intense labeling just above the outer limiting membrane with staining extending into both the nuclear and inner segment layers. In addition, there was a small punctum at the outer segment base where new outer segment discs are formed. At 7 dpi, actin staining in *Pde6g^CreERT2^* experimental retinas looked similar to controls (Fig 6A, 6B). However, at 12 dpi, the inner segment pool of F-actin was greatly depleted, except for a strong band lying just above the outer limiting membrane. This is the location of Muller glia, which have an extensive actin cytoskeleton (Tworig and Feller, 2021) and could be contributing to phalloidin signal in this region. Most actin puncta at the outer segment base were weaker in experimental animals at 12 dpi, but a few strong puncta remained (Fig6A, 6B). The latter puncta likely represent cones as they are located deeper in the inner segment layer and often mark the base of red-green opsin positive outer segments (Fig S6A). To quantitate the intensity of the actin puncta at the base of the connecting cilia in 12 dpi animals, we co-stained phalloidin with an acetylated tubulin antibody, which labels stable microtubules including axonemes (Fig 6C). The number of axonemes in the mutants did not appear to be affected, but the intensity of the actin puncta associated with the axonemes in experimental animals was reduced by about ½ of what was observed in controls (Fig 6D), and the number of puncta were reduced at 12 dpi by ∼1/3 (Fig 6E). Similar results were observed following *iCre75*-driven *Inpp5e* deletion, with inner segment labelling reduced at P15 and largely gone by P21, except for the band above the outer limiting membrane (Fig S5).

**Figure 6.**
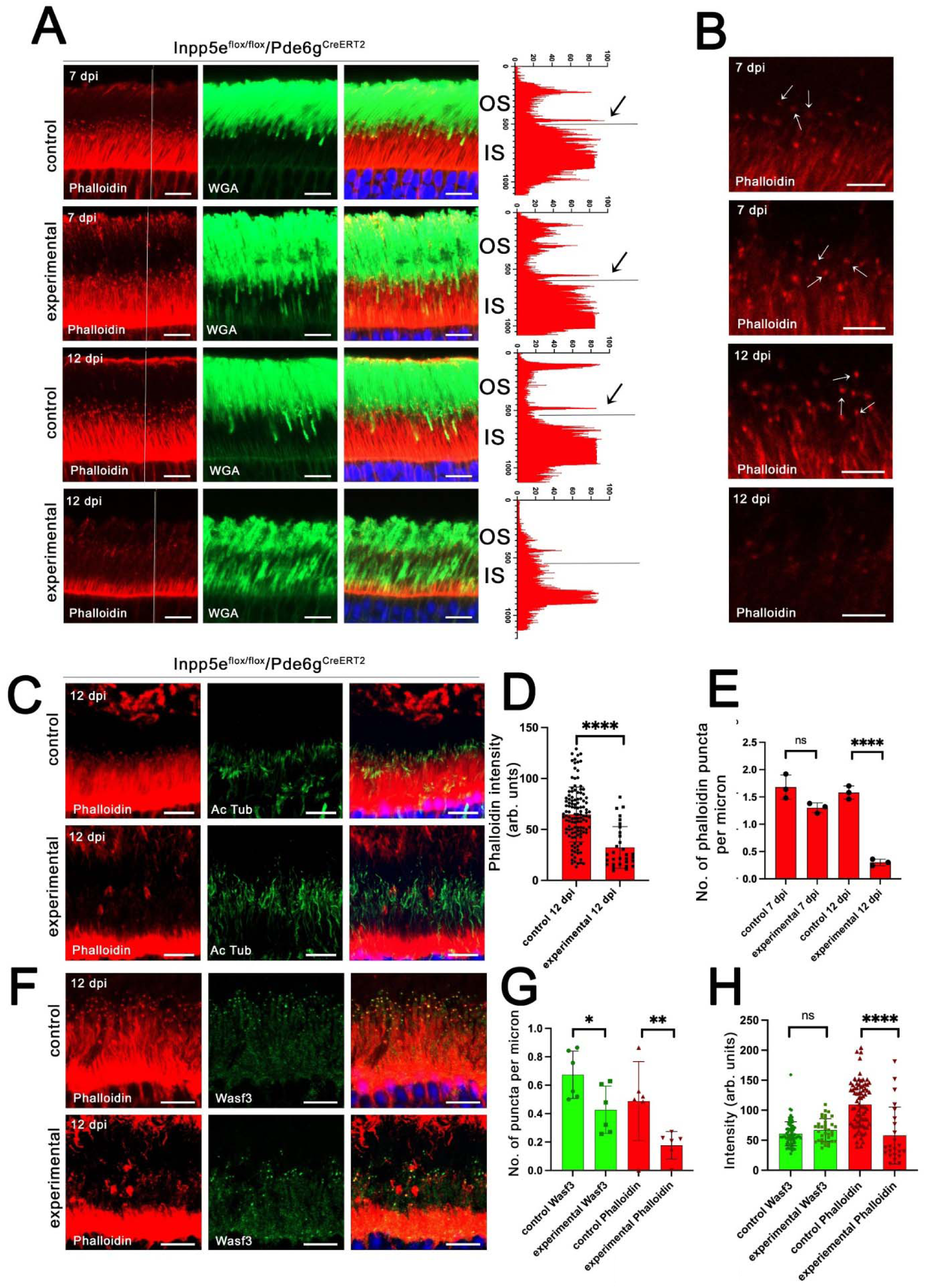
Loss of Inpp5e disrupts the actin cytoskeleton. (A) Confocal images of retinal sections of vehicle- (control) and tamoxifen- (experimental) treated *Inpp5e^flox/flox^/Pde6g^CreERT2/+^*littermates at 7 and 12 day post last injection (dpi) stained with phalloidin (red), and wheat germ agglutinin (WGA, green). Phalloidin intensity along the white line is shown on the right side of the images. Arrows point to signal originating from an actin punctum at the base of an outer segment. Scale bar: 20 µm. Each image is a maximum intensity projection of 5 images taken at 0.7-µm intervals. (B) Enlargements of regions from A showing the actin puncta (arrows) at the junction between the connecting cilium and outer segment. Scale bar: 10 µm. Each image is a maximum intensity projection of 5 images taken at 0.7-µm intervals. (C) Confocal images of retinal sections of vehicle- (control) and tamoxifen- (experimental) treated *Inpp5e^flox/flox^/Pde6g^CreERT2/+^*littermates 12 dpi stained with phalloidin (red), and for acetylated tubulin (Ac Tub, green). Scale bar: 10 µm. Each image is a maximum intensity projection of 20 images taken at 0.7-µm intervals. (D) Fluorescence intensity of phalloidin puncta at the junction of the inner and outer segments in 12 dpi animals. N is 10-40 puncta measured from 3 animals per genotype. Each spot is one measurement. ****p<0.0001, by unpaired t-test. (E) Number of phalloidin puncta per linear µm at the inner-outer segment junction in 7 dpi and 12 dpi for vehicle- (control) and tamoxifen- (experimental) treated *Inpp5e^flox/flox^/Pde6g^CreERT2/+^*littermates. The number of phalloidin puncta were counted from images like in C and divided by the width of the image. N = 3 animals per genotype. **p<0.01, ****p<0.0001, ns not significant by unpaired t-test. (F) Confocal images of retinal sections from vehicle- (control) and tamoxifen- (experimental) treated *Inpp5e^flox/flox^/Pde6g^CreERT2/+^*littermates at 12 days post last injection (dpi) stained with phalloidin (red) and for Wasf3 (green). Scale bars: 10 µm (A), 5 µm (B), 20 µm (C). OS, outer segment; IS, inner segment; ONL, outer nuclear layer; OPL, outer plexiform layer; INL, inner nuclear layer. Each image is a maximum intensity projection of 2 images taken at 0.7-µm intervals. (G) Number of Wasf3 and phalloidin puncta per linear micron at the inner-outer segment junction in 12 dpi animals. The number of Wasf3 and phalloidin puncta were counted from images like in A and divided by the width of the image. N = 4 per genotype. **p<0.01 by unpaired t test. (H) Fluorescence intensity of Wasf3 and phalloidin puncta at the junction of the inner and outer segments in 12 dpi animals. N is 40-50 puncta measured from 3 animals per genotype. Each spot is one measurement. ****p<0.0001, ns, not significant by unpaired t-test.

The WAVE (WASP-family verprolin-homologous protein) complex interacts with Arp2/3 to drive branched actin formation (Takenawa and Suetsugu, 2007). In photoreceptors, the WAVE complex subunit Wasf3 serves as a critical activator of Arp2/3 and is essential for nucleating actin polymerization that drives disc formation at the base of the outer segment (Corral-Serrano et al., 2020; Spencer et al., 2023). To determine if the reduced phalloidin signal in the Inpp5e knockout was due to a failure of Wasf3 to localize properly, we co-stained retinal sections with phalloidin and anti-Wasf3 antibodies (Fig 6F). Fewer phalloidin-positive and fewer Wasf3-positive puncta were observed in experimental animals lacking Inpp5e (Fig 6G).

However, the Wasf3 positive puncta were similar in intensity in both control and experimental animals, while phalloidin signal was reduced in the experimentals (Fig 6H). This suggests that F-actin loss at the base of the outer segment is not a consequence of impairment of actin nucleation mediated by the WAVE complex.

## Discussion

Human patients with INPP5E pathogenic variants develop retinal degeneration (Bielas et al., 2009; Jacoby et al., 2009; Sangermano et al., 2021). Previous work using *Six3-Cre* to delete *Inpp5e* in the developing embryonic eye found that loss of Inpp5e largely blocked the formation of rod and cone outer segments and caused a rapid degeneration of the retina such that almost no rod or cone cells remained at P21. Instead of forming outer segments, the axoneme failed to extend beyond the connecting cilium and the interior was filled with unorganized vesicles rather than membrane discs (Sharif et al., 2021). As this study removed Inpp5e prior to the photoreceptor cells becoming post-mitotic and initiating outer segment development, we sought to explore the role of Inpp5e in later stages of outer segment maturation and maintenance. In our study, we used *iCre75* to delete *Inpp5e* during the elongation phase of outer segment formation and the tamoxifen inducible *Pde6g^CreERT2^* to delete *Inpp5e* from mature photoreceptors. Similar to what was observed with *Six3-Cre*, we found that the loss of *Inpp5e* from rod photoreceptor cells leads to the rapid degeneration of these cells. We found that loss of *Inpp5e* by *iCre75* or *Pde6g^CreERT2^* lead to shorter outer segments that lacked evaginating discs at the base. Our findings are consistent with the observation that no discs formed when *Inpp5e* was deleted by *Six3-Cre*. However, the plasma membrane of the outer segments in the *Six3-Cre* model appeared to have expanded without discs and this was not observed in our models.

The loss of Inpp5e was accompanied by accumulation of rhodopsin and, possibly, other proteins bound by WGA in the cell body, and the presence of extracellular vesicles containing rhodopsin. While extracellular vesicles are occasionally seen in wild type retinas (Lewis et al., 2024; Lewis et al., 2022), their prevalence in our models resembled what is found in rhodopsin mutant models, as well as in other models characterized by rhodopsin mislocalization (Blanks et al., 1982; Blanks and Spee, 1992; Dilan et al., 2018; Gupta et al., 2018; Hagstrom et al., 1999; Heckenlively et al., 1995; Lewis et al., 2024; Lewis et al., 2022; Marszalek et al., 2000; Pazour et al., 2002; Spencer, 2023). Recent work from Lewis and colleagues (Lewis et al., 2024) found that extracellular vesicles released from photoreceptor cells degenerating due to *Ift20* deletion contained a high amount of rhodopsin. Interestingly, these extracellular vesicles had higher concentrations of rhodopsin than was found on the inner segment plasma membrane, suggesting that photoreceptors have a mechanism for concentrating mislocalized rhodopsin before releasing it from the cell. Whether the release of extracellular vesicles serves as a protective mechanism whereby cells discard excess rhodopsin to alleviate stress or the accumulation of vesicles in the extracellular environment is toxic to the retina remains to be identified in future studies [reviewed in (Spencer, 2023)].

Inpp5e was initially identified as a Golgi-associated enzyme (Kong et al., 2000). However, this localization has largely been overlooked in recent studies that focused on the ciliary role of the enzyme in regulating conversion of PI(4,5)P_2_ and PI(3,4,5)P_3_ to PI(4)P and PI(3,4)P_2_ (Dyson et al., 2017) to modulate Hedgehog signaling (Chavez et al., 2015; Constable et al., 2020; Garcia-Gonzalo et al., 2015) and ciliary dissasembly (Phua et al., 2017). In the retina, we found that the loss of the loss of Inpp5e leads to vacuolation and morphological changes in the Golgi apparatus suggesting that the Golgi pool of Inpp5e is likely important to the pathology that results from its loss. In addition, we observed elevated rhodopsin and WGA staining in the inner segment, supporting a role for Inpp5e in the Golgi complex or in trafficking within the inner segment.

Phalloidin staining of the actin cytoskeleton was reduced with *Inpp5e* deletion. Phosphoinositides are well known regulators of actin dynamics. Of relevance to our work, the Inpp5e substrates, PI(4,5)P_2_ and PI(3,4,5)P_3_, are thought to promote actin filament assembly (Hilpela et al., 2004; Janmey et al., 2018). A simple model would suggest that the loss of Inpp5e would elevate these phosphoinositides promoting actin polymerization. However, this was not observed, suggesting a more complicated role for Inpp5e. While the mechanism is unknown, it is possible that indirect effects due to the Golgi dysfunction or trafficking abnormalities drive the observed cytoskeletal defects.

## Experimental Procedures

### Mice

The handling of mice in this study was approved by the institutional animal care and use committees of the UMass Chan School of Medicine. The mice were housed under a 12-hour light and 12-hour dark cycle and had unrestricted access to food and water. Wild type C57BL6/J mice were obtained from Jackson Labs (000664, Bar Harbor ME USA). *Inpp5e^flox/flox^*mice (Dyson et al., 2017) were obtained from Dr. Christina Mitchell at Monash University, Australia.

*Pde6g^CreERT2^* mice (Koch et al., 2015) were obtained from Dr. Stephen Tsang of Columbia University. The *iCre75* mice (Li et al., 2005) were obtained from Jackson Labs (015850, Bar Harbor ME USA). Genotyping was performed by PCR using the primers in Table 1. None of the mice carried the *Pde6b^rd1^* or *Crb1^rd8^* retinal degeneration alleles.

**Table 1.**
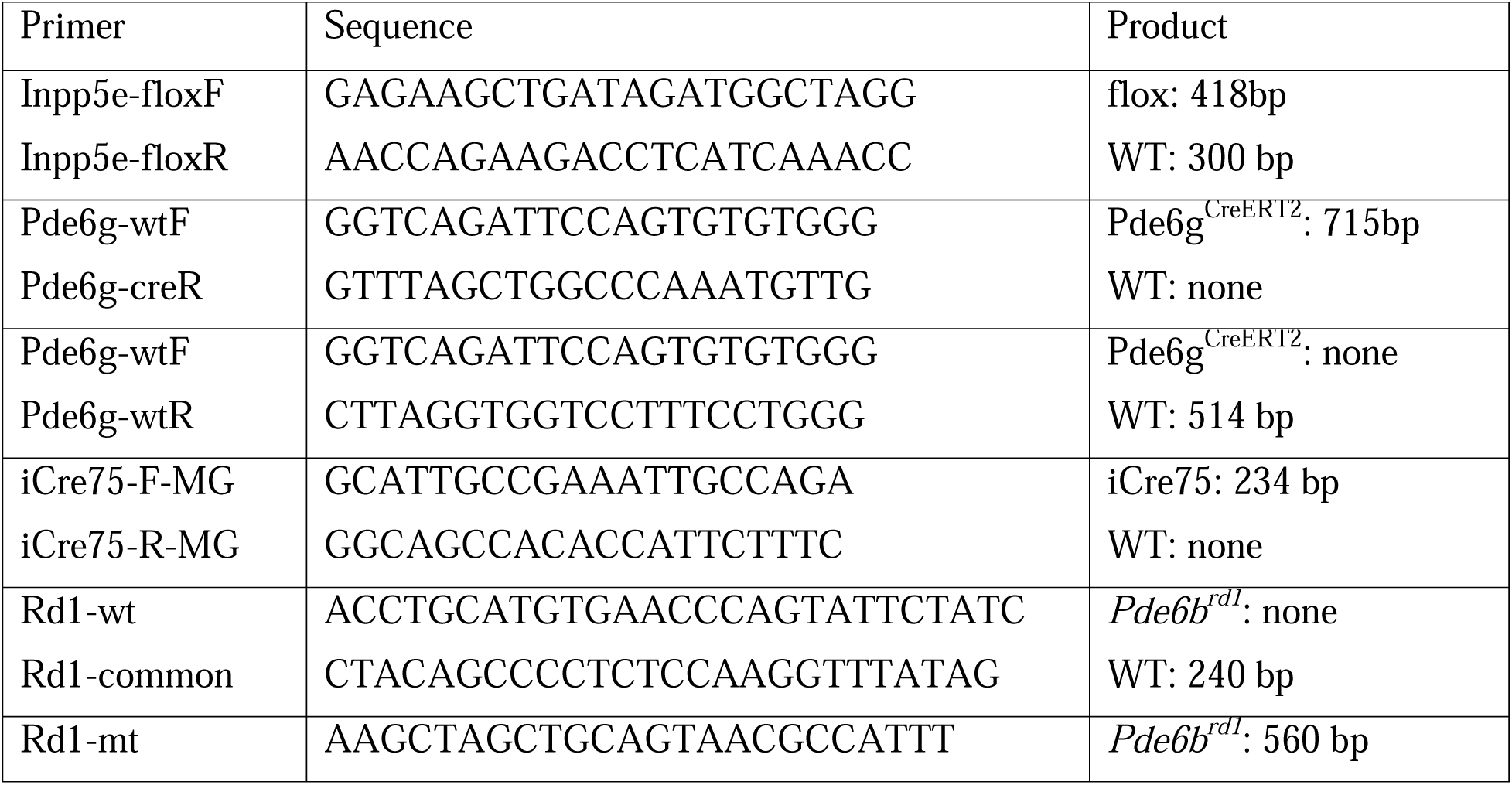
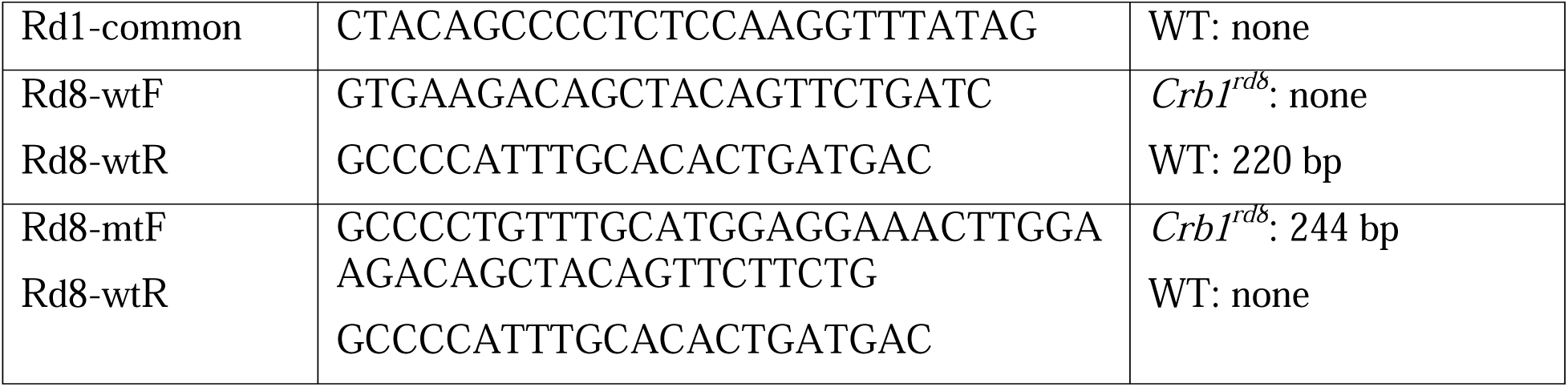
Genotyping primers

For inducible deletion of *Inpp5e* in rod photoreceptor cells by *Pde6g^CreERT2^* 100 microgram of tamoxifen (T5648-1G, Sigma-Aldrich, St Louis MO USA) per gram of body weight was injected intraperitoneally for four consecutive days. Littermates were used as controls.

### Electroretinography

Electroretinogram (ERG) recordings were conducted using a Celeris system (Diagnosys, Lowell MA USA) to evaluate retinal responses under combined dark and light-adapted conditions, including a flicker response. After a 12-hr dark adaptation period, mice were anesthetized via an intraperitoneal injection of a ketamine-xylazine mixture (100 mg/kg and 10 mg/kg, respectively). Pupil dilation was achieved by applying one drop each of phenylephrine (2.5%) and tropicamide (1%) approximately 10 min before recording. Throughout the ERG procedure, animals were maintained on a warming plate to sustain a body temperature of 37°C. ERG electrodes were directly positioned on the eyes and a flicker ERG frequency series was performed with intensities of 0.01 cd·s/m^2^, 0.1 cd·s/m^2^, and 1 cd·s/m^2^, to capture the rod response. Subsequently, a 10-min light adaptation phase was initiated, followed by recording cone impulse responses at intensities of 3 cd·s/m^2^ and 10 cd·s/m^2^, as well as a 10 Hz flicker response. The ERG waveform data were assessed for amplitude and latency of response components, including flicker responses.

### Immunofluorescence Microscopy

Mice were anesthetized by CO_2_ asphyxiation, their eyes removed and placed in 4% paraformaldehyde in Sorensen buffer (100 mM phosphate, pH7.2) overnight at 4°C. After fixation, the eyecups were dissected and embedded in 2.5% low-melt agarose (A3038, Sigma Aldrich). The agarose-embedded eyes were sliced into 100-micrometer thick sections using a vibratome (VT1200S, Leica, Deer Park IL USA) (Lobanova et al., 2010; Spencer et al., 2019).

Sections for immunostaining were blocked in Sorensen buffer containing 7% goat serum and 0.5% Triton X-100 for one hr at room temperature followed by 30 min incubation with antibodies, lectins, and toxins (Table 2). Sections were washed 3 times and incubated with secondary antibodies for 2 hrs at room temperature (Table 2). Following staining, sections were washed three times with PBS and mounted on slides (P36931, Invitrogen, Waltham, MA).

**Table 2.** Antibodies, Lectins, and Toxins

|  | Identifier | Supplier | Concentration |
| --- | --- | --- | --- |
| **Antibody** |  |  |  |
| Inpp5e | 17797-AP | Proteintech,<br>Rosemont IL USA | 1:200 |
| Rhodopsin | Ab98887 (4D2) | Abcam, Cambridge<br>UK | 1:2000 |
| Red/Green Opsin | AB5405 | MilliporeSigma,<br>Burlington, MA USA | 1:2000 |
| Ift27 | 719 | (Keady et al., 2012) | 1:1000 |
| Ift88 | 448 | (Pazour et al., 2002) | 1:1000 |
| Ift140 | 964 | (Jonassen et al., 2012) | 1:1000 |
| Cep164 | 22227-1AP | Santa Cruz Biotech,<br>Santa Cruz CA USA | 1:1000 |
| Prph2 | RDS-CT | (Stuck et al., 2014) | 1:2000 |
| Rom1 |  | (Spencer et al., 2023) | 1:5000 |
| Prom1 | MAB4310 | MilliporeSigma,<br>Burlington, MA USA | 1:100 |
| Stx3 | 15556-1-AP | Proteintech,<br>Rosemont IL USA | 1:2000 |
| Wasf3 | 2806S | Cell Signaling<br>Technology | 1:500 |
| Giantin | 3991 | (Nozawa et al., 2002) | 1:200 |
| Acetylated tubulin | 611-B1<br>(MABT888) | Sigma, St Louis MO<br>USA | 1:2000 |
| Anti-mouse-Alexa488<br>-Alexa594 | A11017<br>A11032 | Invitrogen, Waltham,<br>MA USA | 1:1000 |
| Anti-rabbit-Alexa488<br>-Alexa594 | A11034<br>A21207 | Invitrogen, Waltham,<br>MA USA | 1:1000 |
| **Lectins, toxins, etc.** |  |  |  |
| Wheat germ agglutinin | W11261 | Invitrogen, Waltham, | 1 µg/mL |
| (WGA)-Alexa488 |  | MA USA |  |
| Phalloidin-Alexa594 | A12381 | Invitrogen, Waltham,<br>MA USA | 1:500 |
| 4',6-diamidino-2-phenylindole (DAPI) | D1306 | Invitrogen, Waltham,<br>MA USA | 10 µg/ml |

To prepare cryosections, mouse eyes were fixed in 4% paraformaldehyde in Sorensen buffer for 10 min and then transferred to phosphate buffer without fixative. For cryoprotection, the eyes were sequentially immersed in phosphate buffer supplemented with 10%, 20%, and 30% sucrose for 1 hr, 4 hrs, and overnight, respectively. Subsequently, the eyes were embedded in an optical cutting temperature compound (4583 Tissue-Tek OCT, Sakura, Torrance CA USA) and frozen on dry ice. Using a cryostat, 10-micron sections were cut from the frozen embedded eyes. Sections were blocked in 5% normal goat serum, 0.1% Triton X-100 in Sorensen buffer for 1 hr. Primary and secondary antibodies diluted in blocking buffer were incubated overnight at 4°C and 1 hr at room temperature respectively. After washing nuclei were stained with 10 micrograms/ml 4′,6-diamidino-2-phenylindole (DAPI) for 15 min at room temperature and the tissue mounted with ProLong Gold Antifade Mountant with DAPI (P36931, Vectorlabs, Newark CA USA).

To quantify Golgi morphology, images were analyzed using Fiji/ImageJ software. Ellipticity of Golgi structures was determined by measuring the length of the major and minor axes using the line tool. The major axis was defined as the longest continuous dimension of the Golgi structure. The minor axis was measured as the longest dimension perpendicular to the major axis. Ellipticity was calculated as the ratio of major to minor axis lengths. A minimum of 3 individual Golgi structures were measured from 3 different fields per sample, with 3 biological replicates per condition.

### Transmission Electron Microscopy

Mice were deeply anesthetized and transcardially perfused with 2% paraformaldehyde, 2% glutaraldehyde, 0.05% calcium chloride in 50 mM Mops buffer (pH 7.4) (Ding et al., 2015). Subsequently, the eyes were removed and fixed for an additional 2 hrs in the same buffer at 22°C. After washing in PBS, the eyecups were dissected and embedded in 5% agar (A3038, Sigma-Aldrich) in PBS. 200-micrometer-thick sections were obtained with vibratome (VT1200S, Leica, Deer Park IL USA). The vibratome sections were stained with 1% tannic acid and 1% uranyl acetate (21700 and 22400, Electron Microscopy Sciences, Hatfield PA USA), dehydrated with ethanol, and then infiltrated and embedded in Spurr resin (02680-AB, SPI Supplies, West Chester, PA, USA). 70-nanometer thick sections were cut and placed on copper grids. The sections were counterstained with 2% uranyl acetate and 3.5% lead citrate (19481 and 19314, Ted Pella, Redding CA USA) to enhance contrast. Sections were imaged using either a Philips CM10 electron microscope at 100 kV accelerating voltage with a Gatan TEM CCD camera or a JEOL JEM-1400 electron microscope at 60 kV with a AMT BioSprint camera.

### Statistics

Data groups were analyzed as described in the figure legends using Prism (GraphPad, San Diego CA USA) using unpaired t-tests or ANOVA as described in the figure legends. Differences between groups were considered statistically significant if p < 0.05. Statistical significance is indicated with asterisks (*p=0.01 – 0.05, **p=0.01-0.001, ***p=0.001-0.0001, ****p<0.0001). Error bars are standard deviation (SD).

## Acknowledgements

This work was supported by the National Institutes of Health EY022372 (GJP), EY035525 (VYA), EY005722 (VYA), EY033763 (TRL), E. Matilda Ziegler Foundation for the Blind (WJS) and Unrestricted Awards from Research to Prevent Blindness Inc. (Duke University and SUNY Upstate Medical University). UMass Chan EM core facility resources supported by awards S10 OD025113-01 and S10 OD021580 from the National Center for Research Resources. We thank Dr. Hemant Khanna for initiating this project and obtaining funding, and Dr. Edward Chan (University of Florida) for providing giantin antibodies.

## Author contributions

<u>Mohona Gupta</u>: Conceptualization, Investigation, Data Curation, Visualization, Formal analysis, Writing - Original Draft, Writing - Review & Editing

<u>Tylor R. Lewis</u>: Conceptualization, Investigation, Data Curation, Formal analysis, Writing - Review & Editing

<u>Michael W. Stuck:</u> Conceptualization, Investigation, Formal analysis, Writing - Review & Editing

<u>William J. Spencer</u>: Conceptualization, Investigation, Data Curation, Formal analysis, Writing - Review & Editing

<u>Vadim Y. Arshavsky</u>: Conceptualization, Formal analysis, Writing - Review & Editing, Funding acquisition

<u>Gregory J. Pazour</u>: Conceptualization, Investigation, Formal analysis, Writing - Original Draft, Writing - Review & Editing, Funding acquisition

## Abbreviations

DAPI, 4′,6-diamidino-2-phenylindole; dpi, days post last injection; ERG, Electroretinogram; IFT, Intraflagellar Transport; P, postnatal day; TEM, Transmission electron microscopy;

---

**Figure S1.**
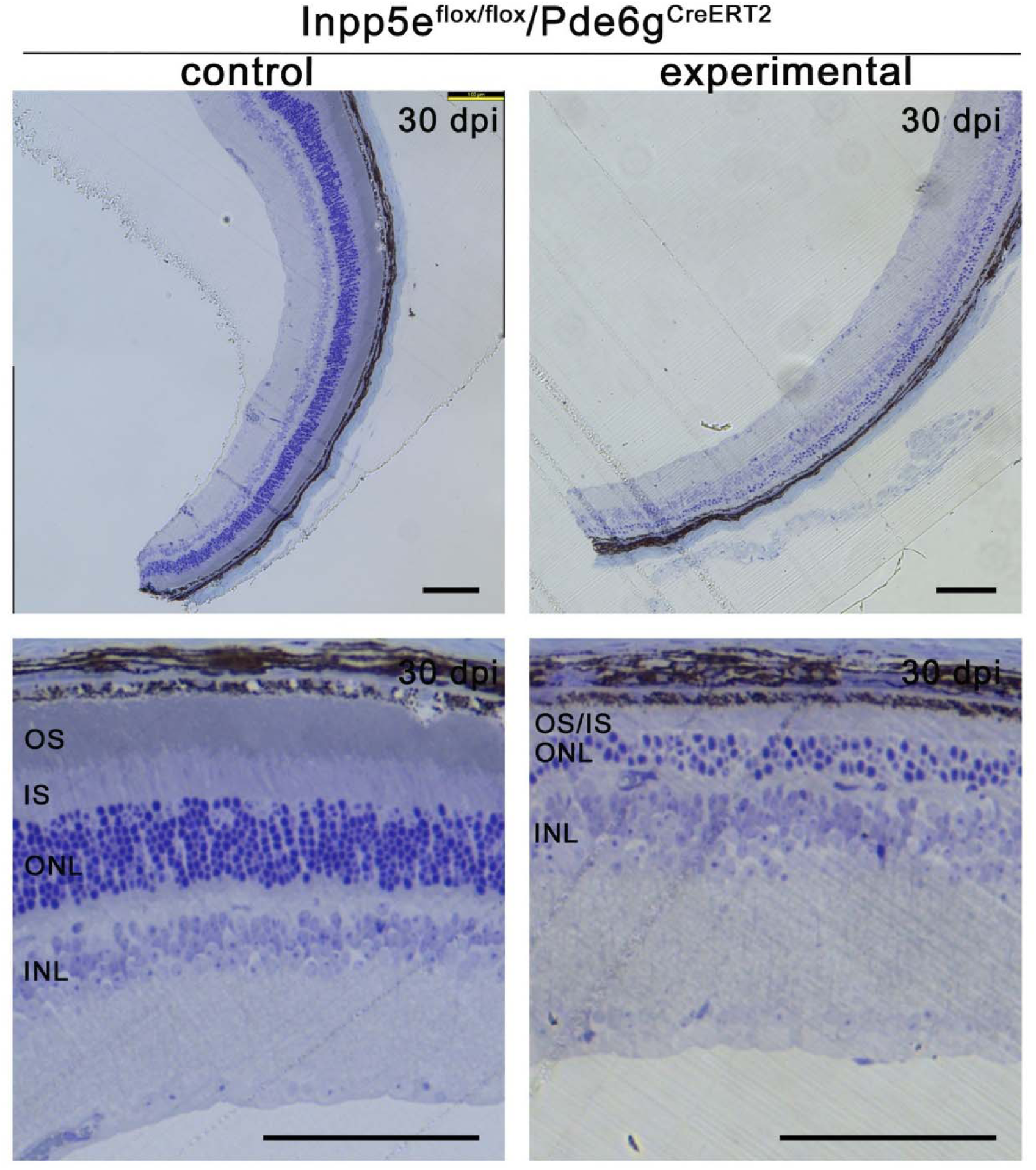
Loss of Inpp5e leads to severe photoreceptor degeneration. Light microscopy of toluidine blue-stained retinal sections from *Inpp5e^flox/flox^/Pde6g^CreERT2^*littermates treated with vehicle (control) or tamoxifen (experimental) and examined at 30 days post last injection. Experimental retinas show severe thinning of the outer nuclear layer (ONL) and near-complete loss of outer segments. OS, outer segment; IS, inner segment; ONL, outer nuclear layer; INL, inner nuclear layer. Scale bar 100 µm.

**Figure S2.**
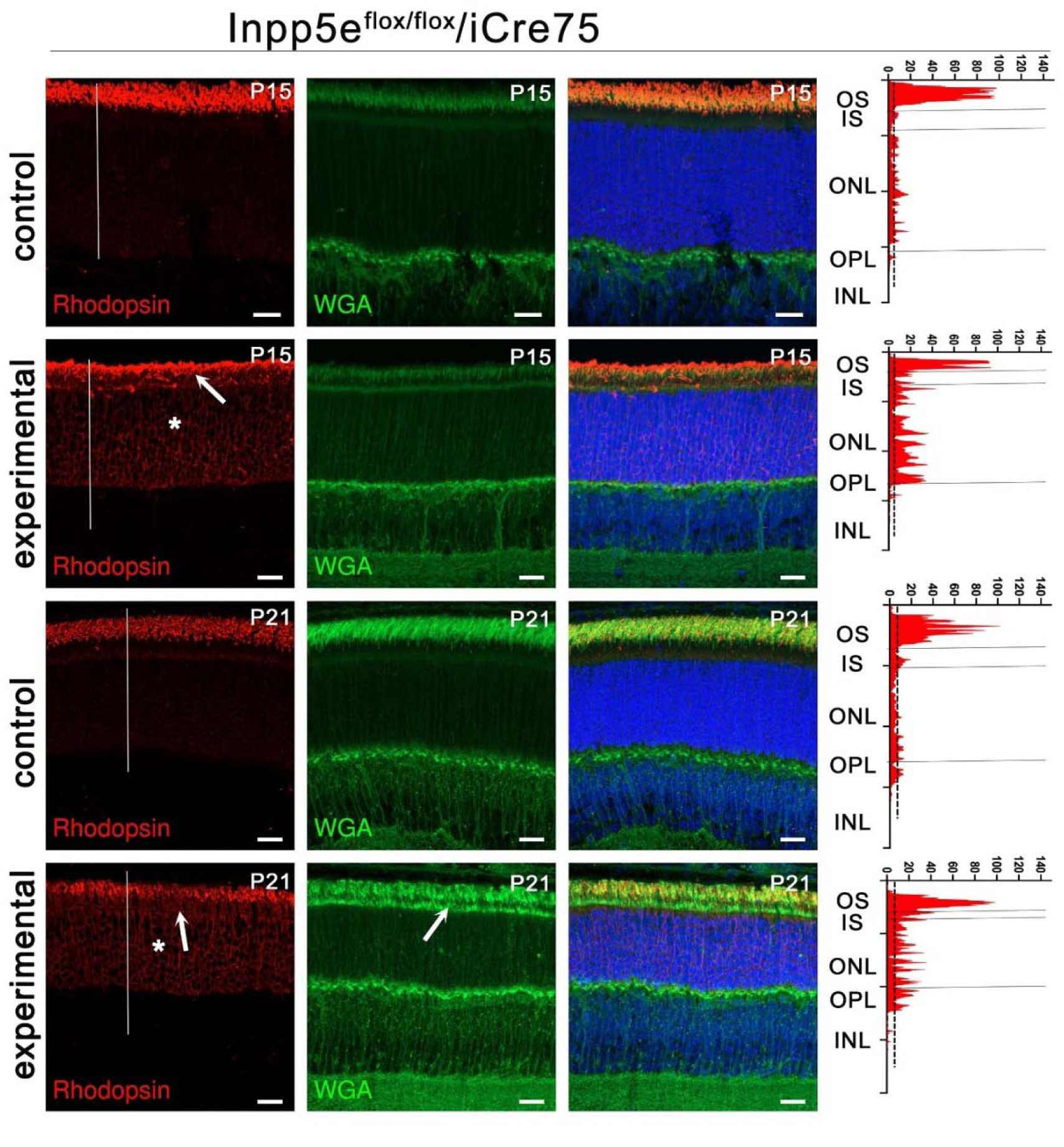
*iCre75*-driven loss of Inpp5e causes rhodopsin mislocalization. (A) Confocal images of retinal sections (agarose embedded) of *Inpp5e^flox/flox^* (control) and *Inpp5e^flox/flox^/iCre75* (experimental) littermates at P15 and P21 immunostained with anti- rhodopsin clone 4D2 (red) and wheat germ agglutinin (WGA, green). Scale bar: 20 µm. Each image is a maximum intensity projection of 20 images taken at 0.7-µm intervals. Rhodopsin intensity along the white line is shown on the right side of the images. Arrows point to mislocalized rhodopsin or WGA in the inner segment. * points to rhodopsin in the nuclear layer. OS, outer segment; IS, inner segment; ONL, outer nuclear layer; OPL, outer plexiform layer; INL, inner nuclear layer.

**Figure S3.**
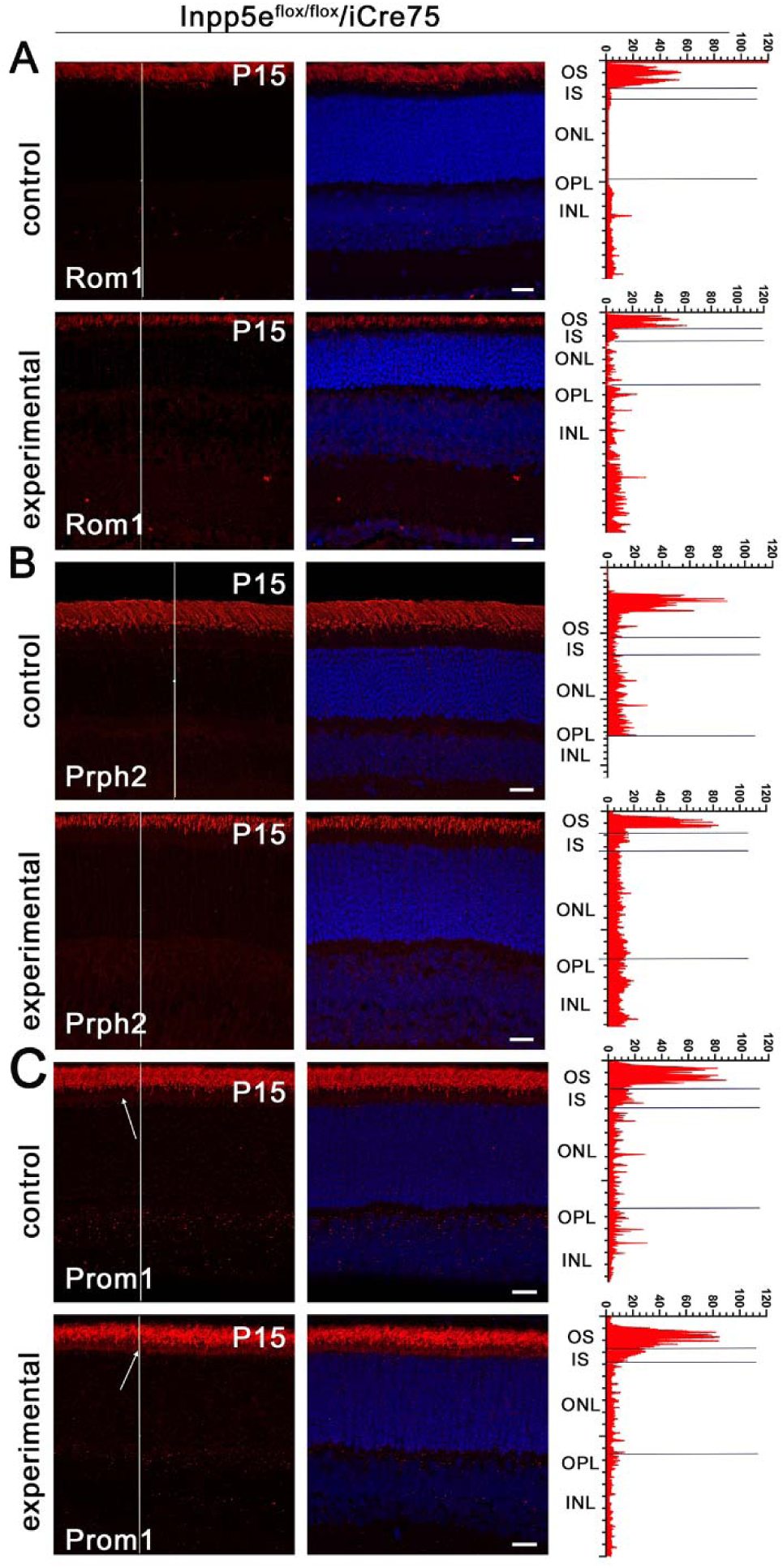
Loss of Inpp5e alters distribution of Prom1 in the *iCre75* cohort. (A-C) Confocal images of retinal sections of *Inpp5e^flox/flox^* (control) and *Inpp5e^flox/flox^/iCre75* (experimental) littermates examined at P15. Sections were stained with DAPI (blue) and for Rom1 (A), Prph2 (B), and Prom1 (C) in red. Note increased Prom1 in the inner segments (arrow) of the experimental animals compared to controls. Scale bars are 20 µm. Each image is a maximum intensity projection of 20 images taken at 0.7-µm intervals. The intensity of the red channel along the white line is shown on the right side of the images. OS, outer segment; IS, inner segment; ONL, outer nuclear layer; OPL, outer plexiform layer; INL, inner nuclear layer.

**Figure S4.**
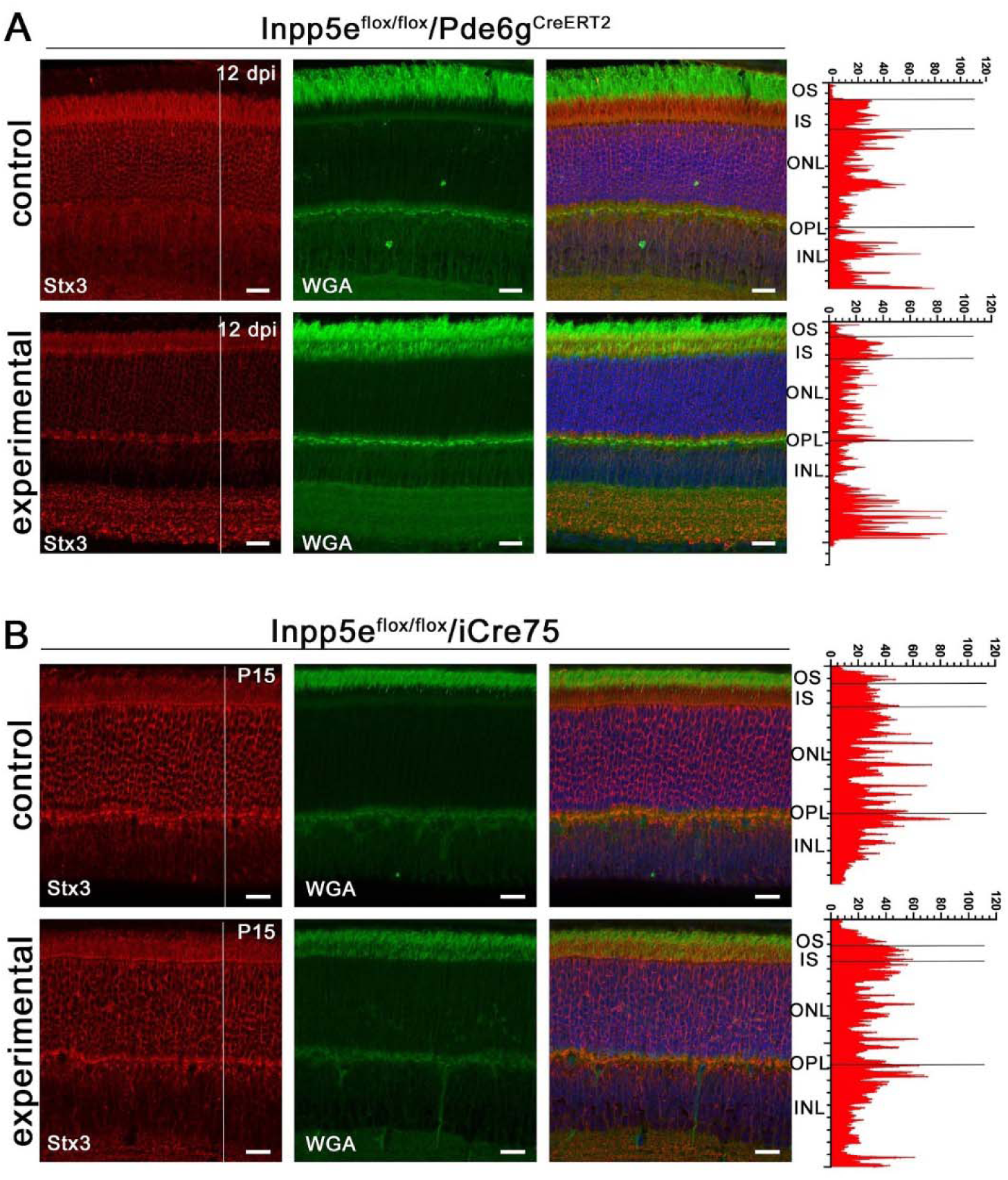
Stx3 is not mislocalized by Inpp5e loss. (A-B) Confocal images of retinal sections of vehicle- (control) and tamoxifen- (experimental) treated *Inpp5e^flox/flox^/Pde6g^CreERT2^* littermates at 12 days post last injection (dpi) (A) or *Inpp5e^flox/flox^* (control) and *Inpp5e^flox/flox^/iCre75* (experimental) littermates examined at P15 (B). Sections were stained with DAPI (blue), wheat germ agglutinin (WGA, green) and syntaxin-3 (Stx3, red). Scale bars are 20 µm. Each image is a maximum intensity projection of 20 images taken at 0.7-µm intervals. The intensity of the red channel along the white line is shown on the right side of the images. OS, outer segment; IS, inner segment; ONL, outer nuclear layer; OPL, outer plexiform layer; INL, inner nuclear layer.

**Figure S5.**
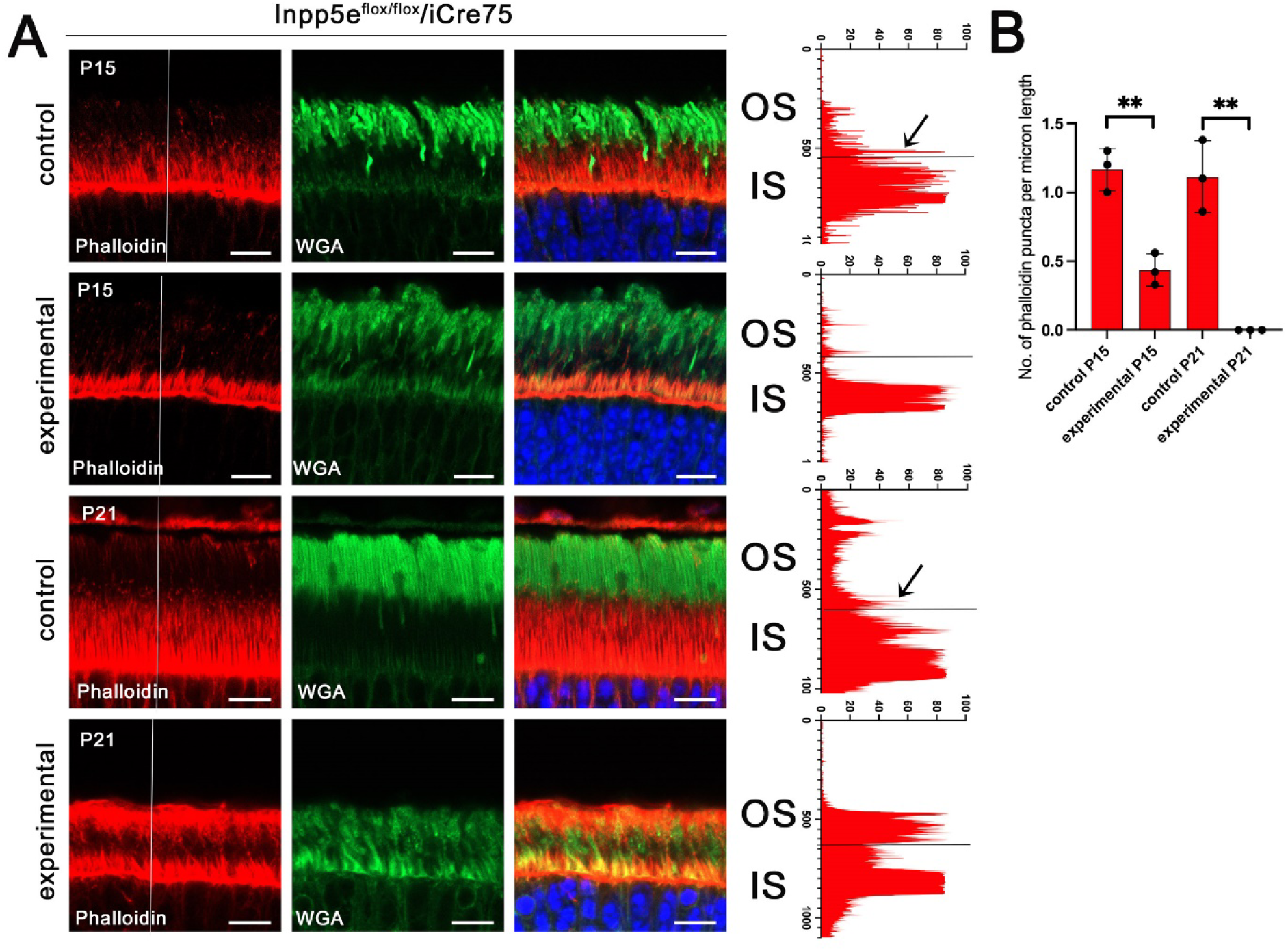
*iCre75*-driven loss of Inpp5e disrupts the actin cytoskeleton. (A) Confocal images of retinal sections of *Inpp5e^flox/flox^* (control) and *Inpp5e^flox/flox^/iCre75* (experimental) littermates at P15 and P21 stained with phalloidin (red), and wheat germ agglutinin (WGA, green). Phalloidin intensity along the white line is shown on the right side of the images. Arrows point to signal originating from an actin punctum at the base of an outer segment. Scale bar: 20 µm. Each image is a maximum intensity projection of 2 images taken at 0.7-µm intervals. (B) Number of phalloidin puncta per linear µm at the inner-outer segment junction in P15 and P21 animals. The number of phalloidin puncta were counted from images like in A and divided by the width of the image. N = 3 animals per genotype. **p<0.01, by unpaired t-test.

**Figure S6.**
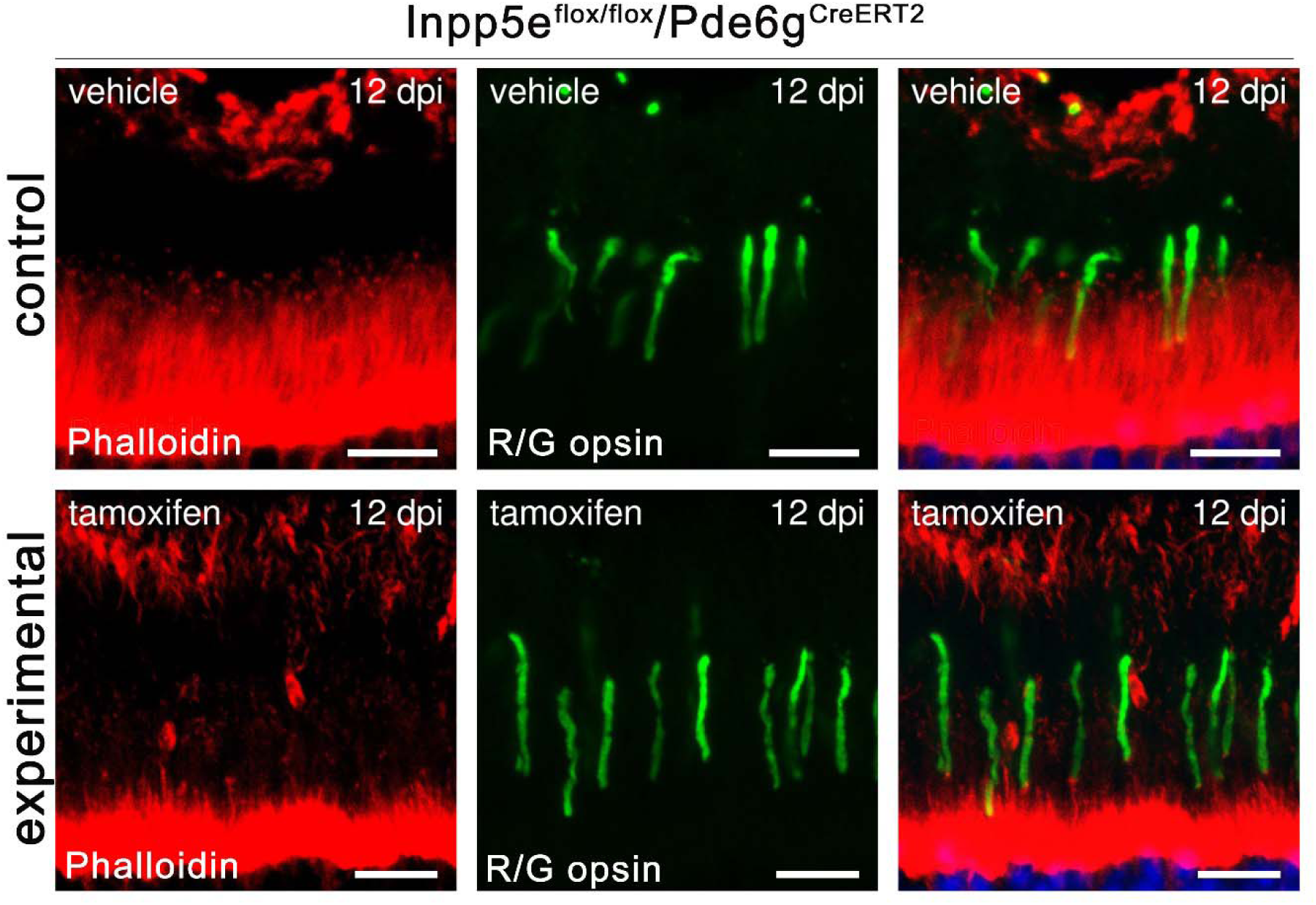
Actin puncta remain at the base of cones. Confocal images of retinal sections of vehicle- (control) and tamoxifen- (experimental) treated *Inpp5e^flox/flox^/Pde6g^CreERT2/+^* littermates at 12 days post last injection (dpi) stained with phalloidin (red), and red/green opsin (R/G opsin, green). Scale bars:10 µm. Each image is a maximum intensity projection of 8 images taken at 0.7-µm intervals.

## References

1. Bielas, S.L., J.L. Silhavy, F. Brancati, M.V. Kisseleva, L. Al-Gazali, L. Sztriha, R.A. Bayoumi, M.S. Zaki, A. Abdel-Aleem, R.O. Rosti, H. Kayserili, D. Swistun, L.C. Scott, E. Bertini, E. Boltshauser, E. Fazzi, L. Travaglini, S.J. Field, S. Gayral, M. Jacoby, S. Schurmans, B. Dallapiccola, P.W. Majerus, E.M. Valente, and J.G. Gleeson. 2009. Mutations in INPP5E, encoding inositol polyphosphate-5-phosphatase E, link phosphatidyl inositol signaling to the ciliopathies. Nat Genet. 41:1032–1036. PMID: 19668216. 10.1038/ng.423. PMC2746682

2. Blanks, J.C., R.J. Mullen, and M.M. LaVail. 1982. Retinal degeneration in the pcd cerebellar mutant mouse. II. Electron microscopic analysis. J Comp Neurol. 212:231–246. PMID: 7153375. 10.1002/cne.902120303.

3. Blanks, J.C., and C. Spee. 1992. Retinal degeneration in the pcd/pcd mutant mouse: accumulation of spherules in the interphotoreceptor space. Exp Eye Res. 54:637–644. PMID: 1623950. 10.1016/0014-4835(92)90019-o.

4. Campos, B., L. Zeng, P.H. Daotrong, V. Eckstein, A. Unterberg, H. Mairbaurl, and C. Herold-Mende. 2011. Expression and regulation of AC133 and CD133 in glioblastoma. Glia. 59:1974–1986. PMID: 21901757. 10.1002/glia.21239.

5. Chavez, M., S. Ena, J. Van Sande, A. de Kerchove d’Exaerde, S. Schurmans, and S.N. Schiffmann. 2015. Modulation of Ciliary Phosphoinositide Content Regulates Trafficking and Sonic Hedgehog Signaling Output. Dev Cell. 34:338–350. PMID: 26190144. 10.1016/j.devcel.2015.06.016.

6. Constable, S., A.B. Long, K.A. Floyd, S. Schurmans, and T. Caspary. 2020. The ciliary phosphatidylinositol phosphatase Inpp5e plays positive and negative regulatory roles in Shh signaling. Development. 147. PMID: 31964774. 10.1242/dev.183301. PMC7033722

7. Corral-Serrano, J.C., I.J.C. Lamers, J. van Reeuwijk, L. Duijkers, A.D.M. Hoogendoorn, A. Yildirim, N. Argyrou, R.A.A. Ruigrok, S.J.F. Letteboer, R. Butcher, M.D. van Essen, S. Sakami, S.E.C. van Beersum, K. Palczewski, M.E. Cheetham, Q. Liu, K. Boldt, U. Wolfrum, M. Ueffing, A. Garanto, R. Roepman, and R.W.J. Collin. 2020. PCARE and WASF3 regulate ciliary F-actin assembly that is required for the initiation of photoreceptor outer segment disk formation. Proc Natl Acad Sci U S A. 117:9922–9931. PMID: 32312818. 10.1073/pnas.1903125117. PMC7211956

8. Datta, P., C. Allamargot, J.S. Hudson, E.K. Andersen, S. Bhattarai, A.V. Drack, V.C. Sheffield, and S. Seo. 2015. Accumulation of non-outer segment proteins in the outer segment underlies photoreceptor degeneration in Bardet-Biedl syndrome. Proc Natl Acad Sci U S A. 112:E4400–4409. PMID: 26216965. 10.1073/pnas.1510111112. PMC4538681

9. Dewees, S.I., R. Vargova, K.R. Hardin, R.E. Turn, S. Devi, J. Linnert, U. Wolfrum, T. Caspary, M. Elias, and R.A. Kahn. 2022. Phylogenetic profiling and cellular analyses of ARL16 reveal roles in traffic of IFT140 and INPP5E. Mol Biol Cell. 33:ar33. PMID: 35196065. 10.1091/mbc.E21-10-0509-T.PMC9250359

10. Dilan, T.L., R.K. Singh, T. Saravanan, A. Moye, A.F.X. Goldberg, P. Stoilov, and V. Ramamurthy. 2018. Bardet-Biedl syndrome-8 (BBS8) protein is crucial for the development of outer segments in photoreceptor neurons. Hum Mol Genet. 27:283–294. PMID: 29126234. 10.1093/hmg/ddx399. PMC5886228

11. Ding, J.D., R.Y. Salinas, and V.Y. Arshavsky. 2015. Discs of mammalian rod photoreceptors form through the membrane evagination mechanism. J Cell Biol. 211:495–502. PMID: 26527746. 10.1083/jcb.201508093. PMC4639867

12. Dyson, J.M., S.E. Conduit, S.J. Feeney, S. Hakim, T. DiTommaso, A.J. Fulcher, A. Sriratana, G. Ramm, K.A. Horan, R. Gurung, C. Wicking, I. Smyth, and C.A. Mitchell. 2017. INPP5E regulates phosphoinositide-dependent cilia transition zone function. J Cell Biol. 216:247–263. PMID: 27998989. 10.1083/jcb.201511055. PMC5223597

13. Furuta, Y., O. Lagutin, B.L. Hogan, and G.C. Oliver. 2000. Retina- and ventral forebrain-specific Cre recombinase activity in transgenic mice. Genesis. 26:130–132. PMID: 10686607.

14. Garcia-Gonzalo, F.R., S.C. Phua, E.C. Roberson, G. Garcia, 3rd, M. Abedin, S. Schurmans, T. Inoue, and J.F. Reiter. 2015. Phosphoinositides Regulate Ciliary Protein Trafficking to Modulate Hedgehog Signaling. Dev Cell. 34:400-409. PMID: 26305592. 10.1016/j.devcel.2015.08.001. PMC4557815

15. Gupta, M., and G.J. Pazour. 2023. Intraflagellar transport: A critical player in photoreceptor development and the pathogenesis of retinal degenerative diseases. Cytoskeleton (Hoboken). PMID: 38140908. 10.1002/cm.21823.

16. Gupta, P.R., N. Pendse, S.H. Greenwald, M. Leon, Q. Liu, E.A. Pierce, and K.M. Bujakowska. 2018. Ift172 conditional knock-out mice exhibit rapid retinal degeneration and protein trafficking defects. Hum Mol Genet. 27:2012–2024. PMID: 29659833. 10.1093/hmg/ddy109. PMC5961092

17. Hagstrom, S.A., M. Duyao, M.A. North, and T. Li. 1999. Retinal degeneration in tulp1-/- mice: vesicular accumulation in the interphotoreceptor matrix. Invest Ophthalmol Vis Sci. 40:2795–2802. PMID: 10549638.

18. Hampshire, D.J., M. Ayub, K. Springell, E. Roberts, H. Jafri, Y. Rashid, J. Bond, J.H. Riley, and C.G. Woods. 2006. MORM syndrome (mental retardation, truncal obesity, retinal dystrophy and micropenis), a new autosomal recessive disorder, links to 9q34. Eur J Hum Genet. 14:543–548. PMID: 16493448. 10.1038/sj.ejhg.5201577.

19. Hasegawa, J., R. Iwamoto, T. Otomo, A. Nezu, M. Hamasaki, and T. Yoshimori. 2016. Autophagosome-lysosome fusion in neurons requires INPP5E, a protein associated with Joubert syndrome. EMBO J. 35:1853–1867. PMID: 27340123. 10.15252/embj.201593148. PMC5007553

20. Heckenlively, J.R., B. Chang, L.C. Erway, C. Peng, N.L. Hawes, G.S. Hageman, and T.H. Roderick. 1995. Mouse model for Usher syndrome: linkage mapping suggests homology to Usher type I reported at human chromosome 11p15. Proc Natl Acad Sci U S A. 92:11100–11104. PMID: 7479945. 10.1073/pnas.92.24.11100. PMC40579

21. Hilpela, P., M.K. Vartiainen, and P. Lappalainen. 2004. Regulation of the actin cytoskeleton by PI(4,5)P2 and PI(3,4,5)P3. Curr Top Microbiol Immunol. 282:117–163. PMID: 14594216. 10.1007/978-3-642-18805-3_5.

22. Hsu, Y., J.E. Garrison, G. Kim, A.R. Schmitz, C.C. Searby, Q. Zhang, P. Datta, D.Y. Nishimura, S. Seo, and V.C. Sheffield. 2017. BBSome function is required for both the morphogenesis and maintenance of the photoreceptor outer segment. PLoS Genet. 13:e1007057. PMID: 29049287. 10.1371/journal.pgen.1007057. PMC5663628

23. Jacoby, M., J.J. Cox, S. Gayral, D.J. Hampshire, M. Ayub, M. Blockmans, E. Pernot, M.V. Kisseleva, P. Compere, S.N. Schiffmann, F. Gergely, J.H. Riley, D. Perez-Morga, C.G. Woods, and S. Schurmans. 2009. INPP5E mutations cause primary cilium signaling defects, ciliary instability and ciliopathies in human and mouse. Nat Genet. 41:1027–1031. PMID: 19668215.10.1038/ng.427.

24. Janmey, P.A., R. Bucki, and R. Radhakrishnan. 2018. Regulation of actin assembly by PI(4,5)P2 and other inositol phospholipids: An update on possible mechanisms. Biochem Biophys Res Commun. 506:307–314. PMID: 30139519. 10.1016/j.bbrc.2018.07.155. PMC6269227

25. Keady, B.T., R. Samtani, K. Tobita, M. Tsuchya, J.T. San Agustin, J.A. Follit, J.A. Jonassen, R. Subramanian, C.W. Lo, and G.J. Pazour. 2012. IFT25 links the signal-dependent movement of Hedgehog components to intraflagellar transport. Dev Cell. 22:940–951. PMID: 22595669. 10.1016/j.devcel.2012.04.009. PMC3366633

26. Kisseleva, M.V., M.P. Wilson, and P.W. Majerus. 2000. The isolation and characterization of a cDNA encoding phospholipid-specific inositol polyphosphate 5-phosphatase. J Biol Chem. 275:20110–20116. PMID: 10764818. 10.1074/jbc.M910119199.

27. Koch, S.F., Y.T. Tsai, J.K. Duong, W.H. Wu, C.W. Hsu, W.P. Wu, L. Bonet-Ponce, C.S. Lin, and S.H. Tsang. 2015. Halting progressive neurodegeneration in advanced retinitis pigmentosa. J Clin Invest. 125:3704–3713. PMID: 26301813. 10.1172/JCI82462. PMC4588299

28. Kong, A.M., C.J. Speed, C.J. O’Malley, M.J. Layton, T. Meehan, K.L. Loveland, S. Cheema, L.M. Ooms, and C.A. Mitchell. 2000. Cloning and characterization of a 72-kDa inositol-polyphosphate 5-phosphatase localized to the Golgi network. J Biol Chem. 275:24052–24064. PMID: 10806194. 10.1074/jbc.M000874200.

29. LaVail, M.M. 1973. Kinetics of rod outer segment renewal in the developing mouse retina. J Cell Biol. 58:650–661. PMID: 4747920. 10.1083/jcb.58.3.650. PMC2109077

30. Lewis, T.R., C.M. Castillo, N.V. Klementieva, Y. Hsu, Y. Hao, W.J. Spencer, A.V. Drack, G.J. Pazour, and V.Y. Arshavsky. 2024. Contribution of intraflagellar transport to compartmentalization and maintenance of the photoreceptor cell. Proc Natl Acad Sci U S A. 121:e2408551121. PMID: 39145934. 10.1073/pnas.2408551121.

31. Lewis, T.R., S. Phan, K.Y. Kim, I. Jha, C.M. Castillo, J.D. Ding, B.S. Sajdak, D.K. Merriman, M.H. Ellisman, and V.Y. Arshavsky. 2022. Microvesicle release from inner segments of healthy photoreceptors is a conserved phenomenon in mammalian species. Dis Model Mech. 15. PMID: 36420970. 10.1242/dmm.049871. PMC9796728

32. Li, S., D. Chen, Y. Sauve, J. McCandless, Y.J. Chen, and C.K. Chen. 2005. Rhodopsin-iCre transgenic mouse line for Cre-mediated rod-specific gene targeting. Genesis. 41:73–80. PMID: 15682388. 10.1002/gene.20097.

33. Lobanova, E.S., R. Herrmann, S. Finkelstein, B. Reidel, N.P. Skiba, W.T. Deng, R. Jo, E.R. Weiss, W.W. Hauswirth, and V.Y. Arshavsky. 2010. Mechanistic basis for the failure of cone transducin to translocate: why cones are never blinded by light. J Neurosci. 30:6815–6824. PMID: 20484624. 10.1523/JNEUROSCI.0613-10.2010. PMC2883257

34. Marszalek, J.R., X. Liu, E.A. Roberts, D. Chui, J.D. Marth, D.S. Williams, and L.S. Goldstein. 2000. Genetic evidence for selective transport of opsin and arrestin by kinesin-II in mammalian photoreceptors. Cell. 102:175–187. PMID: 10943838. 10.1016/s0092-8674(00)00023-4.

35. Nozawa, K., C.A. Casiano, J.C. Hamel, C. Molinaro, M.J. Fritzler, and E.K. Chan. 2002. Fragmentation of Golgi complex and Golgi autoantigens during apoptosis and necrosis. Arthritis Res. 4:R3. PMID: 12106502. 10.1186/ar422. PMC125295

36. Obata, S., and J. Usukura. 1992. Morphogenesis of the photoreceptor outer segment during postnatal development in the mouse (BALB/c) retina. Cell Tissue Res. 269:39–48. PMID: 1423483. 10.1007/BF00384724.

37. Papermaster, D.S., B.G. Schneider, D. DeFoe, and J.C. Besharse. 1986. Biosynthesis and vectorial transport of opsin on vesicles in retinal rod photoreceptors. J Histochem Cytochem. 34:5–16. PMID: 2934469. 10.1177/34.1.2934469.

38. Pazour, G.J., S.A. Baker, J.A. Deane, D.G. Cole, B.L. Dickert, J.L. Rosenbaum, G.B. Witman, and J.C. Besharse. 2002. The intraflagellar transport protein, IFT88, is essential for vertebrate photoreceptor assembly and maintenance. *J Cell Biol*. 157:103-113. PMID: 11916979. 10.1083/jcb.200107108. PMC2173265

39. Pearring, J.N., R.Y. Salinas, S.A. Baker, and V.Y. Arshavsky. 2013. Protein sorting, targeting and trafficking in photoreceptor cells. Prog Retin Eye Res. 36:24–51. PMID: 23562855. 10.1016/j.preteyeres.2013.03.002. PMC3759535

40. Phua, S.C., S. Chiba, M. Suzuki, E. Su, E.C. Roberson, G.V. Pusapati, S. Schurmans, M. Setou, R. Rohatgi, J.F. Reiter, K. Ikegami, and T. Inoue. 2017. Dynamic Remodeling of Membrane Composition Drives Cell Cycle through Primary Cilia Excision. Cell. 168:264–279 e215. PMID: 28086093. 10.1016/j.cell.2016.12.032. PMC5660509

41. Sangermano, R., I. Deitch, V.G. Peter, R. Ba-Abbad, E.M. Place, E. Zampaglione, N.E. Wagner, A.B. Fulton, L. Coutinho-Santos, B. Rosin, V. Dunet, A. AlTalbishi, E. Banin, A.B. Sousa, M. Neves, A. Larson, M. Quinodoz, M. Michaelides, T. Ben-Yosef, E.A. Pierce, C. Rivolta, A.R. Webster, G. Arno, D. Sharon, R.M. Huckfeldt, and K.M. Bujakowska. 2021. Broadening INPP5E phenotypic spectrum: detection of rare variants in syndromic and non-syndromic IRD. NPJ Genom Med. 6:53. PMID: 34188062. 10.1038/s41525-021-00214-8. PMC8242099

42. Sharif, A.S., C.D. Gerstner, M.A. Cady, V.Y. Arshavsky, C. Mitchell, G. Ying, J.M. Frederick, and W. Baehr. 2021. Deletion of the phosphatase INPP5E in the murine retina impairs photoreceptor axoneme formation and prevents disc morphogenesis. J Biol Chem. 296:100529. PMID: 33711342. 10.1016/j.jbc.2021.100529. PMC8047226

43. Spencer, W.J. 2023. Extracellular vesicles highlight many cases of photoreceptor degeneration. Front Mol Neurosci. 16:1182573. PMID: 37273908. 10.3389/fnmol.2023.1182573. PMC10233141

44. Spencer, W.J., T.R. Lewis, J.N. Pearring, and V.Y. Arshavsky. 2020. Photoreceptor Discs: Built Like Ectosomes. Trends Cell Biol. 30:904–915. PMID: 32900570. 10.1016/j.tcb.2020.08.005. PMC7584774

45. Spencer, W.J., T.R. Lewis, S. Phan, M.A. Cady, E.O. Serebrovskaya, N.F. Schneider, K.Y. Kim, L.A. Cameron, N.P. Skiba, M.H. Ellisman, and V.Y. Arshavsky. 2019. Photoreceptor disc membranes are formed through an Arp2/3-dependent lamellipodium-like mechanism. Proc Natl Acad Sci U S A. 116:27043–27052. PMID: 31843915. 10.1073/pnas.1913518117. PMC6936530

46. Spencer, W.J., N.F. Schneider, T.R. Lewis, C.M. Castillo, N.P. Skiba, and V.Y. Arshavsky. 2023. The WAVE complex drives the morphogenesis of the photoreceptor outer segment cilium. Proc Natl Acad Sci U S A. 120:e2215011120. PMID: 36917665. 10.1073/pnas.2215011120. PMC10041111

47. Stuck, M.W., S.M. Conley, and M.I. Naash. 2014. The Y141C knockin mutation in RDS leads to complex phenotypes in the mouse. Hum Mol Genet. 23:6260–6274. PMID: 25001182. 10.1093/hmg/ddu345. PMC4222364

48. Stuck, M.W., S.M. Conley, and M.I. Naash. 2016. PRPH2/RDS and ROM-1: Historical context, current views and future considerations. Prog Retin Eye Res. 52:47–63. PMID: 26773759. 10.1016/j.preteyeres.2015.12.002. PMC4842342

49. Sung, C.H., and A.W. Tai. 2000. Rhodopsin trafficking and its role in retinal dystrophies. Int Rev Cytol. 195:215–267. PMID: 10603577. 10.1016/s0074-7696(08)62706-0.

50. Takenawa, T., and S. Suetsugu. 2007. The WASP-WAVE protein network: connecting the membrane to the cytoskeleton. Nat Rev Mol Cell Biol. 8:37–48. PMID: 17183359. 10.1038/nrm2069.

51. Tian, G., P. Ropelewski, I. Nemet, R. Lee, K.H. Lodowski, and Y. Imanishi. 2014. An unconventional secretory pathway mediates the cilia targeting of peripherin/rds. J Neurosci. 34:992–1006. PMID: 24431457. 10.1523/JNEUROSCI.3437-13.2014. PMC3891973

52. Travaglini, L., F. Brancati, J. Silhavy, M. Iannicelli, E. Nickerson, N. Elkhartoufi, E. Scott, E. Spencer, S. Gabriel, S. Thomas, B. Ben-Zeev, E. Bertini, E. Boltshauser, M. Chaouch, M.R. Cilio, M.M. de Jong, H. Kayserili, G. Ogur, A. Poretti, S. Signorini, G. Uziel, M.S. Zaki, J.S.G. International, C. Johnson, T. Attie-Bitach, J.G. Gleeson, and E.M. Valente. 2013. Phenotypic spectrum and prevalence of INPP5E mutations in Joubert syndrome and related disorders. Eur J Hum Genet. 21:1074–1078. PMID: 23386033. 10.1038/ejhg.2012.305. PMC3778343

53. Tworig, J.M., and M.B. Feller. 2021. Muller Glia in Retinal Development: From Specification to Circuit Integration. Front Neural Circuits. 15:815923. PMID: 35185477. 10.3389/fncir.2021.815923. PMC8856507

54. Van De Weghe, J.C., A. Gomez, and D. Doherty. 2022. The Joubert-Meckel-Nephronophthisis Spectrum of Ciliopathies. Annu Rev Genomics Hum Genet. 23:301–329. PMID: 35655331. 10.1146/annurev-genom-121321-093528. PMC9437135

55. Yang, Z., Y. Chen, C. Lillo, J. Chien, Z. Yu, M. Michaelides, M. Klein, K.A. Howes, Y. Li, Y. Kaminoh, H. Chen, C. Zhao, Y. Chen, Y.T. Al-Sheikh, G. Karan, D. Corbeil, P. Escher, S. Kamaya, C. Li, S. Johnson, J.M. Frederick, Y. Zhao, C. Wang, D.J. Cameron, W.B. Huttner, D.F. Schorderet, F.L. Munier, A.T. Moore, D.G. Birch, W. Baehr, D.M. Hunt, D.S. Williams, and K. Zhang. 2008. Mutant prominin 1 found in patients with macular degeneration disrupts photoreceptor disk morphogenesis in mice. J Clin Invest. 118:2908–2916. PMID: 18654668. 10.1172/JCI35891. PMC2483685

56. Young, R.W. 1967. The renewal of photoreceptor cell outer segments. J Cell Biol. 33:61–72. PMID: 6033942. 10.1083/jcb.33.1.61. PMC2107286

