## Supplemental Figures for "Inpp5e Is Critical for Photoreceptor Outer Segment Maintenance"

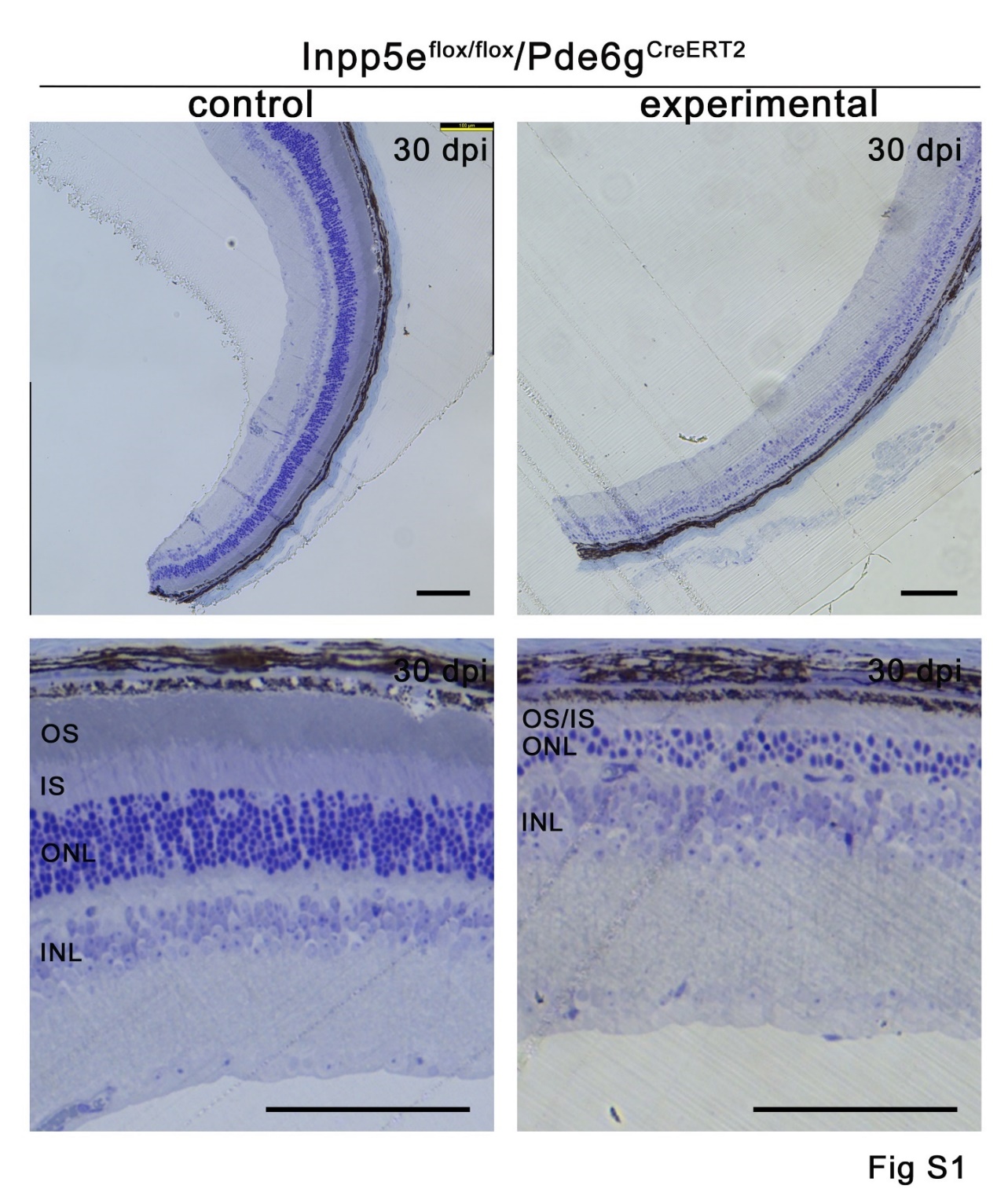

### Figure S1. Loss of Inpp5e leads to severe photoreceptor degeneration.

Light microscopy of toluidine blue-stained retinal sections from *Inpp5e^flox/flox^/Pde6g^CreERT2^* littermates treated with vehicle (control) or tamoxifen (experimental) and examined at 30 days post last injection. Experimental retinas show severe thinning of the outer nuclear layer (ONL) and near-complete loss of outer segments. OS, outer segment; IS, inner segment; ONL, outer nuclear layer; INL, inner nuclear layer. Scale bar 100 µm.

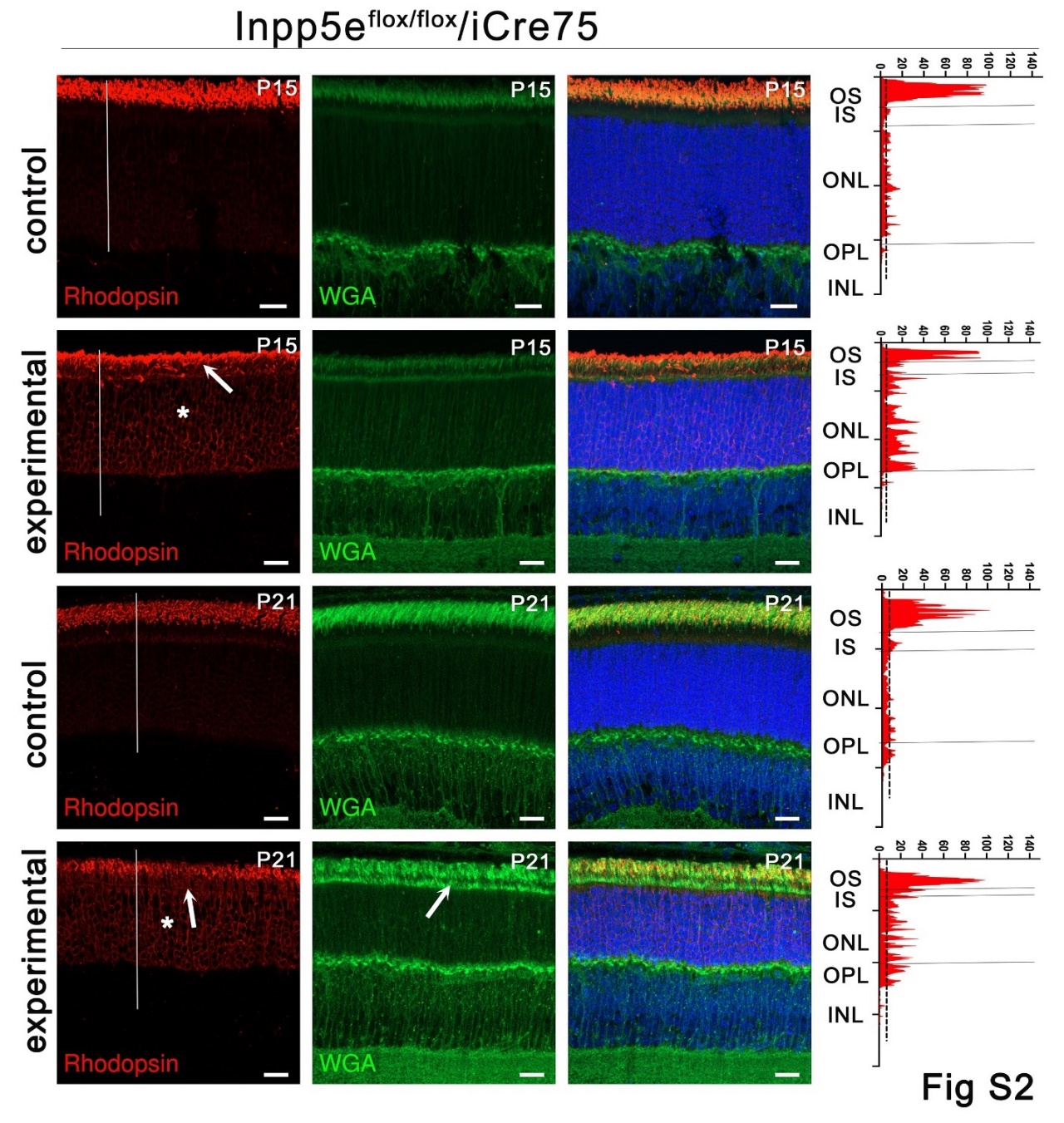

### Figure S2. *iCre75*-driven loss of Inpp5e causes rhodopsin mislocalization.

(A) Confocal images of retinal sections (agarose embedded) of *Inpp5e^flox/flox^* (control) and *Inpp5e^flox/flox^/iCre75* (experimental) littermates at P15 and P21 immunostained with anti-rhodopsin clone 4D2 (red) and wheat germ agglutinin (WGA, green). Scale bar: 20 µm. Each image is a maximum intensity projection of 20 images taken at 0.7-µm intervals. Rhodopsin intensity along the white line is shown on the right side of the images. Arrows point to mislocalized rhodopsin or WGA in the inner segment. * points to rhodopsin in the nuclear layer. OS, outer segment; IS, inner segment; ONL, outer nuclear layer; OPL, outer plexiform layer; INL, inner nuclear layer.

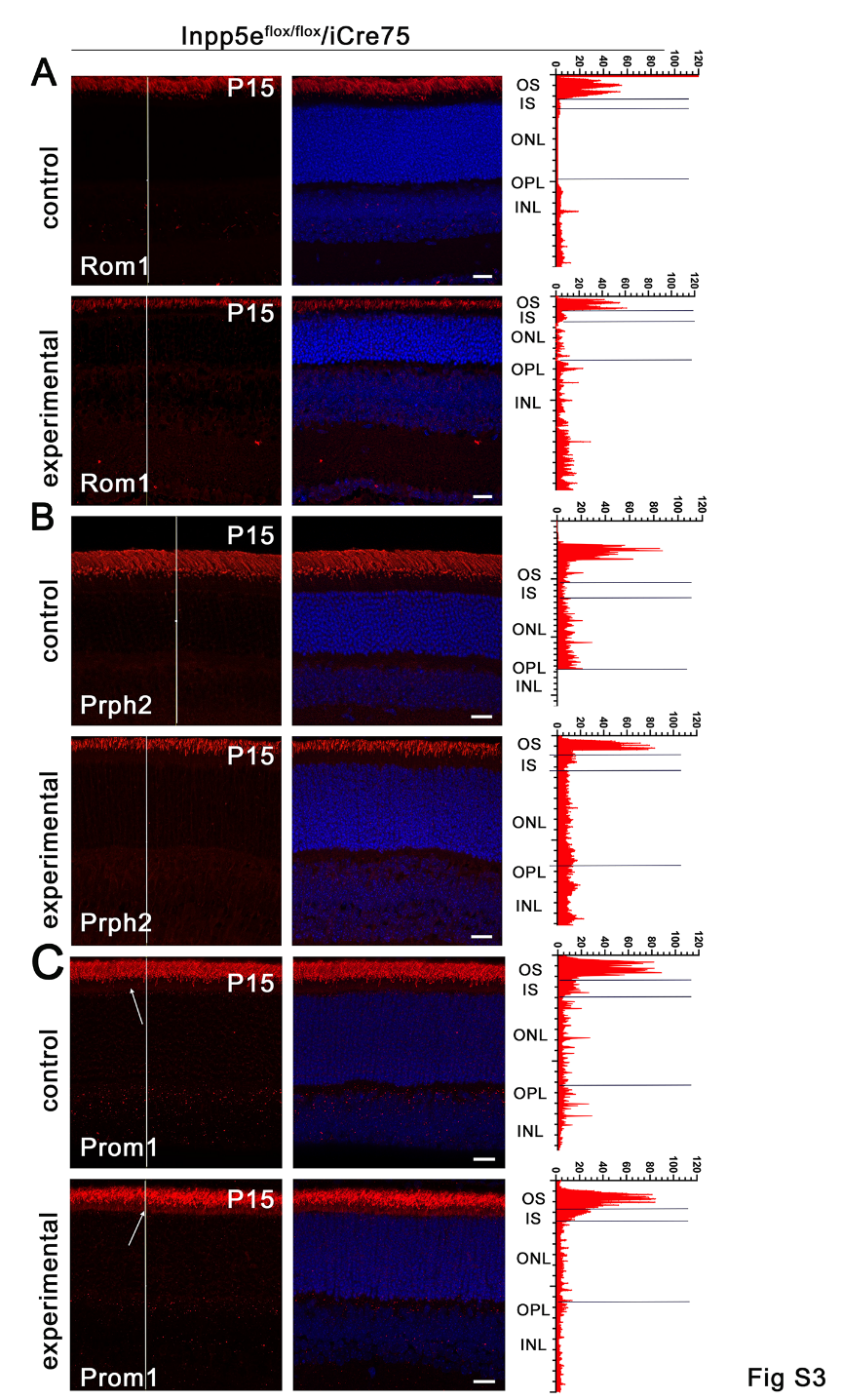

### Figure S3. Loss of Inpp5e alters distribution of Prom1 in the *iCre75* cohort.

(A-C) Confocal images of retinal sections of *Inpp5e^flox/flox^* (control) and *Inpp5e^flox/flox^/iCre75* (experimental) littermates examined at P15. Sections were stained with DAPI (blue) and for Rom1 (A), Prph2 (B), and Prom1 (C) in red. Note increased Prom1 in the inner segments (arrow) of the experimental animals compared to controls. Scale bars are 20 µm. Each image is a maximum intensity projection of 20 images taken at 0.7-µm intervals. The intensity of the red channel along the white line is shown on the right side of the images. OS, outer segment; IS, inner segment; ONL, outer nuclear layer; OPL, outer plexiform layer; INL, inner nuclear layer.

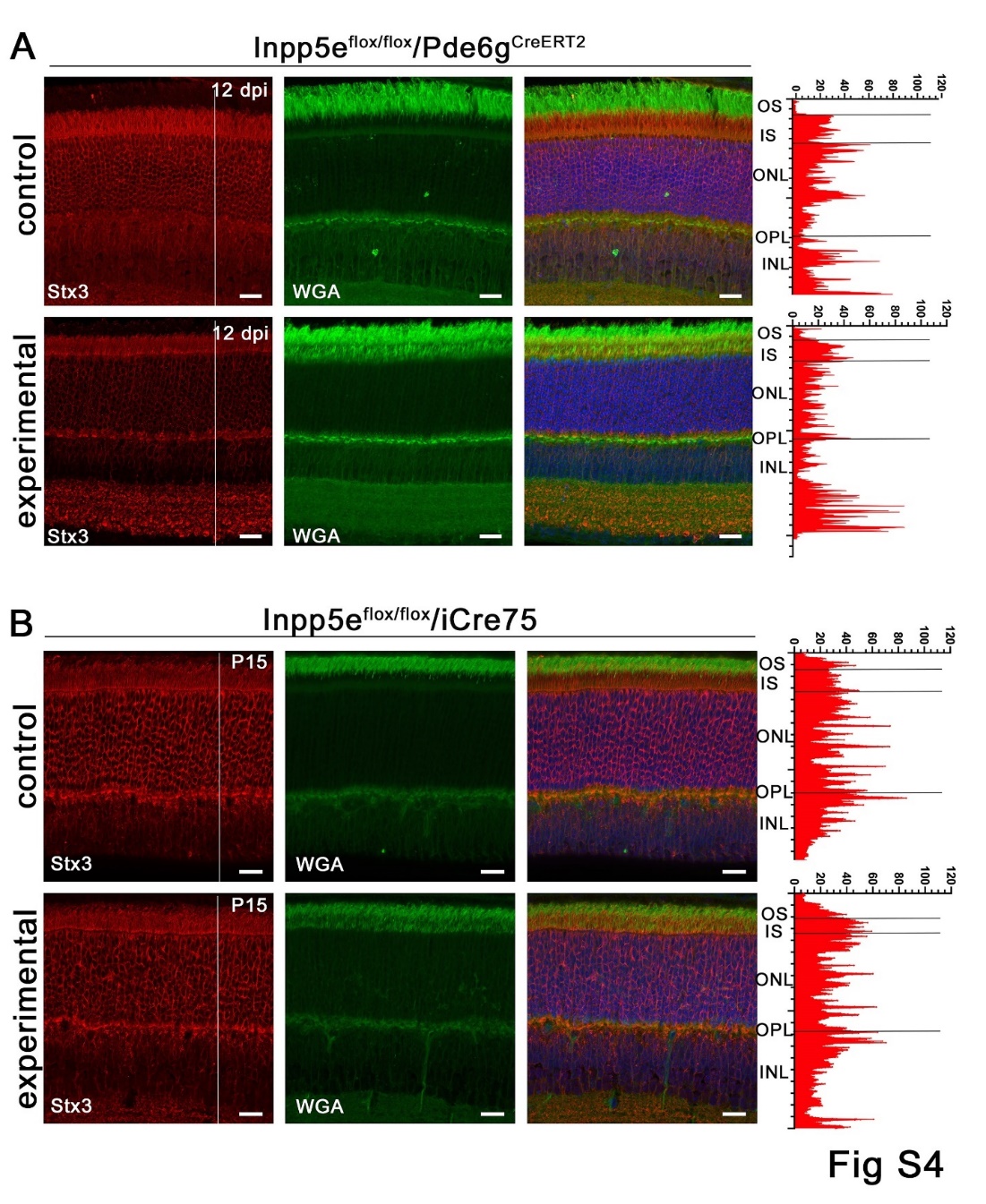

### Figure S4. Stx3 is not mislocalized by Inpp5e loss.

(A-B) Confocal images of retinal sections of vehicle- (control) and tamoxifen- (experimental) treated *Inpp5e^flox/flox^/Pde6g^CreERT2^* littermates at 12 days post last injection (dpi) (A) or *Inpp5e^flox/flox^* (control) and *Inpp5e^flox/flox^/iCre75* (experimental) littermates examined at P15 (B). Sections were stained with DAPI (blue), wheat germ agglutinin (WGA, green) and syntaxin-3 (Stx3, red). Scale bars are 20 µm. Each image is a maximum intensity projection of 20 images taken at 0.7-µm intervals. The intensity of the red channel along the white line is shown on the right side of the images. OS, outer segment; IS, inner segment; ONL, outer nuclear layer; OPL, outer plexiform layer; INL, inner nuclear layer.

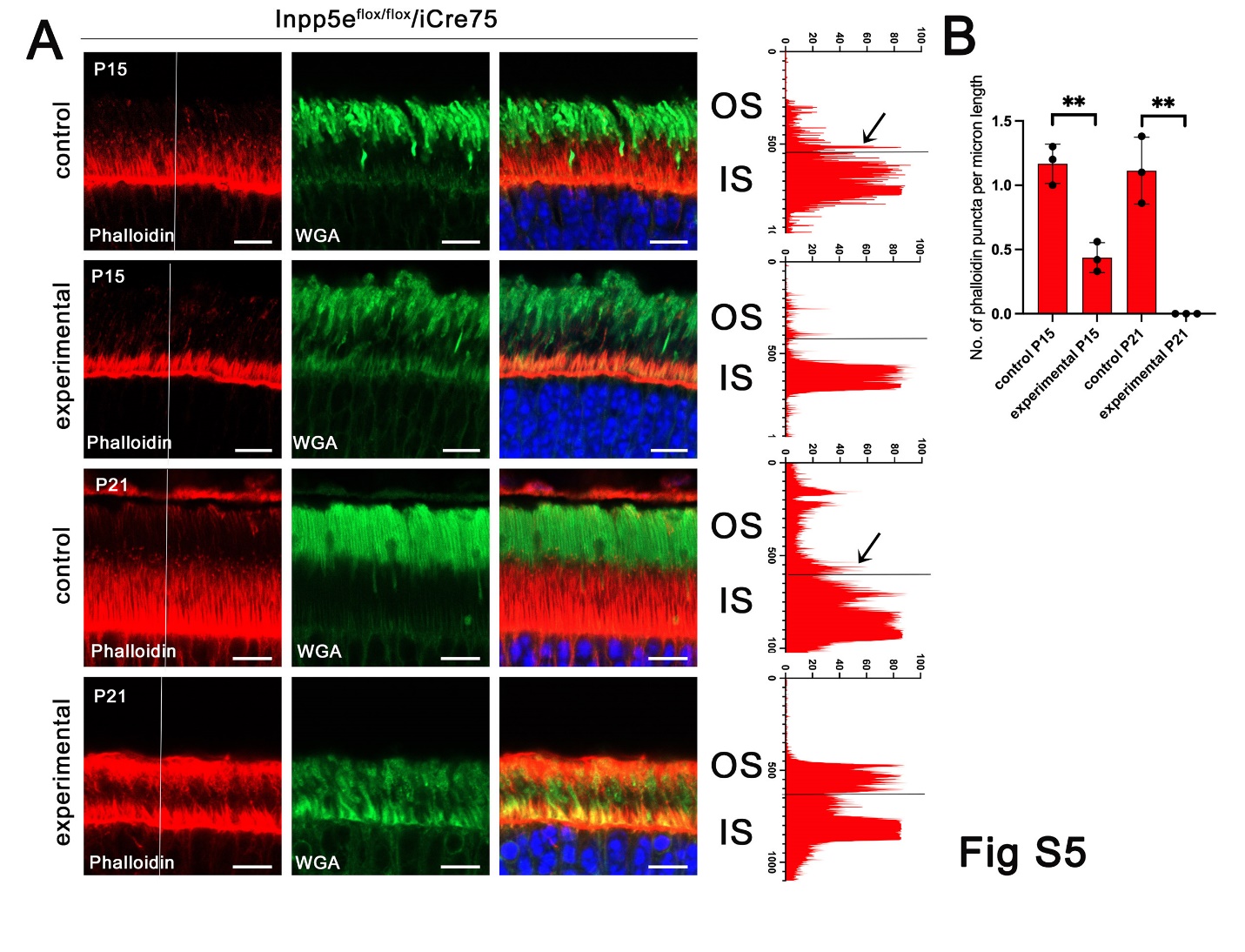

### Figure S5. *iCre75*-driven loss of Inpp5e disrupts the actin cytoskeleton.

(A) Confocal images of retinal sections of *Inpp5e^flox/flox^* (control) and *Inpp5e^flox/flox^/iCre75* (experimental) littermates at P15 and P21 stained with phalloidin (red), and wheat germ agglutinin (WGA, green). Phalloidin intensity along the white line is shown on the right side of the images. Arrows point to signal originating from an actin punctum at the base of an outer segment. Scale bar: 20 µm. Each image is a maximum intensity projection of 2 images taken at 0.7-µm intervals.

(B) Number of phalloidin puncta per linear µm at the inner-outer segment junction in P15 and P21 animals. The number of phalloidin puncta were counted from images like in A and divided by the width of the image. N = 3 animals per genotype. **p<0.01, by unpaired t-test.

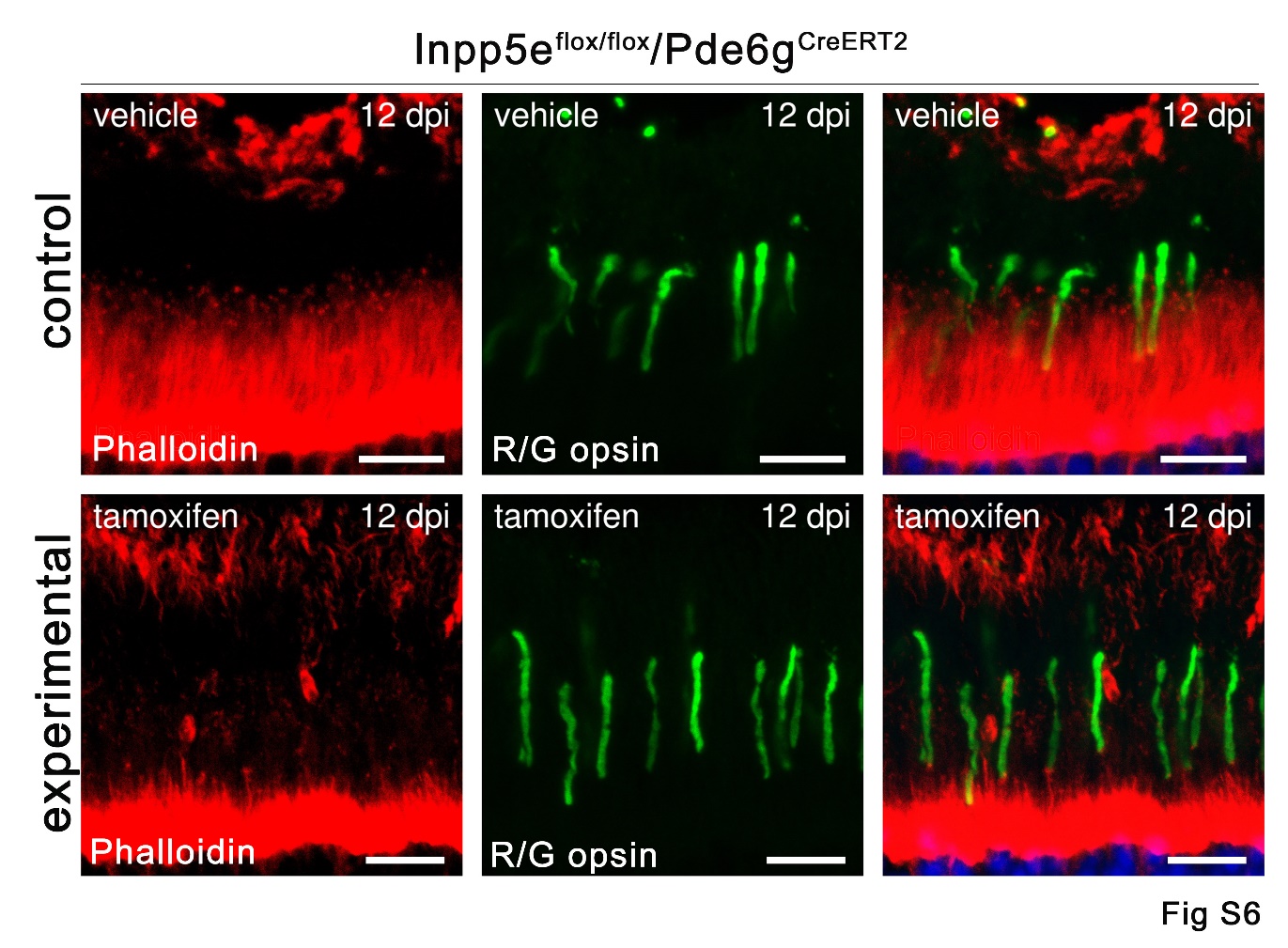

### Figure S6. Actin puncta remain at the base of cones**.**

Confocal images of retinal sections of vehicle- (control) and tamoxifen- (experimental) treated *Inpp5e^flox/flox^/Pde6g^CreERT2/+^* littermates at 12 days post last injection (dpi) stained with phalloidin (red), and red/green opsin (R/G opsin, green). Scale bars:10 µm. Each image is a maximum intensity projection of 8 images taken at 0.7-µm intervals.
